# Inhibition of cathepsin B blocks amyloidogenesis in the mouse models of neurological lysosomal diseases mucopolysaccharidosis type IIIC and sialidosis

**DOI:** 10.1101/2025.01.20.633731

**Authors:** Gustavo M. Viana, Xuefang Pan, Shuxian Fan, TianMeng Xu, Alexandra Wyatt, Alexey V. Pshezhetsky

## Abstract

Neuronal accumulation of amyloid aggregates is a hallmark of brain pathology in neurological lysosomal storage diseases (LSDs) including mucopolysaccharidoses (MPS), however, the molecular mechanism underlaying this pathology has not been understood. We demonstrate that elevated lysosomal cathepsin B (CTSB) levels and CTSB leakage to the cytoplasm triggers amyloidogenesis in two neurological LSDs. CTSB levels were elevated 3-5-fold in the cortices of mouse models of MPS IIIC (*Hgsnat-Geo* and *Hgsnat^P304L^*) and sialidosis (*Neu1^ΔEx3^*), as well as in cortical samples of MPS I, IIIA, IIIC and IIID patients. CTSB was found in the cytoplasm of pyramidal layer IV-V cortical neurons containing Thioflavin-S-positive, β-amyloid-positive aggregates consistent with pro-senile phenotype. In contrast, CTSB-deficient MPS IIIC (*Hgsnat^P304L^/Ctsb^-/-^*) mice as well as *Hgsnat^P304L^* and *Neu1^ΔEx3^* mice chronically treated with irreversible brain-penetrable CTSB inhibitor, E64, showed a drastic reduction of neuronal Thioflavin-S-positive/APP-positive deposits. Neurons of *Hgsnat^P304L^/Ctsb^-/-^* mice and E64-treated *Hgsnat^P304L^* mice also showed reduced levels of P62/LC3-positive puncta, G_M2_ ganglioside and misfolded subunit C of mitochondrial ATP synthase (SCMAS) consistent with restored autophagy. E64 treatment also rescued hyperactivity and reduced anxiety in *Hgsnat^P304L^* mice implying that CTSB may become a novel pharmacological target for MPS III and similar LSDs.

## Introduction

About two-thirds of patients affected with lysosomal storage diseases (LSDs), inherited metabolic disorders caused by lysosomal dysfunction, display neurological symptoms [1]. Although the severity of CNS pathology and specific pathological signs can be different for different LSDs, several pathological cascades were found to be common. This includes neuroimmune response, manifesting with population of the brain by activated macrophages/microglia and astrocytes expressing pro-inflammatory cytokines, neuronal death affecting essential brain areas such as cortex, hippocampus and cerebellum, loss of myelin and neuronal dysfunction affecting synaptic transmission and synaptogenesis (reviewed in the ref. [2]). Several common biomarkers of CNS pathology in neurological LSDs, most notably amyloidogenesis and accumulation of neuronal misfolded protein aggregates, neurofibrillary tangles and lipofuscin bodies, are also shared with adult neurogenerative disorders, such as Parkinson disease, Alzheimer’s disease, frontotemporal dementia, dementia with Lewy bodies and others. Besides, prevalence of common neurogenerative diseases is often associated with the deficiency or haploinsufficiency of lysosomal hydrolases such as glucocerebrosidase GBA, acid sphingomyelinase, and others suggesting existence of a shared pathophysiological mechanisms with LSDs such as defects in the autophagic pathway, essential for removal of misfolded and aggregated proteins or damaged organelles [3–7].

Intraneuronal aggregates of beta amyloid peptides have been observed in multiple classes of LSDs with different types of primary biochemical defects and storage materials. In particular, it has been detected in neurological mucopolysaccharidoses (MPS) caused by deficiency of lysosomal enzymes involved in degradation of heparan sulfate (HS) [8]. In the mouse models of MPS type III (Sanfilippo disease), caused by deficiency of 4 lysosomal enzymes specifically involved in HS catabolism, amyloid aggregates were predominantly found in the pyramidal cortical neurons, that also accumulated misfolded mitochondrial proteins (mainly subunit C of mitochondrial ATP synthase, SCMAS), phosphorylated Tau and secondary metabolites such as cholesterol and simple gangliosides (e.g. G_M2_ and G_M3_) [9–14]. This has been proposed to cause an intracellular metabolic imbalance and, consequently, loss of neuronal viability [12,15,16]. The neurons containing β-amyloid deposits were also strongly positive for the presence of P62- positive and LC3-positive puncta, the markers of impaired autophagy, however, the exact mechanistic link between the two phenomena has not been established and requires further investigation (reviewed in ref. [17]). In particular, inhibition of autophagy and lysosomal genes expression corrected several pathological features in MPS III cells and MPS IIIB model [18] as well as in MPS II mice [19], however induction of autophagy and lysosomal genes expression ameliorated phenotype of MPS IIIA mice [20]. Notably, accumulation of β-amyloid aggregates was also described in the mouse model of a neurological LSD sialidosis, caused by genetic deficiency of Neuraminidase 1 (NEU1) [21].

The authors speculated that NEU1 deficiency caused oversialylation of β-amyloid peptides and their increased secretion from the cell leading to formation of amyloid plaques [21].

The internalization and subsequent processing of the amyloid precursor protein (APP) by β-secretase 1 (BACE-1) and γ-secretase have been recognized as the main mechanism underlying generation of β-amyloid peptides in the AD [22–24]. However, other proteases and, in particular, lysosomal cathepsin B (CTSB) also play a role in amyloidogenic APP processing [25–29]. Importantly, CTSB was also found to be an essential component of the inflammasome, required for inflammatory activation of microglia [30–32], and one of the key enzymes in regulation of autophagic flux [33]. Previously, we have shown in the cortical neurons of the MPS I mouse model that increased CTSB levels coincided coincided with amyloidogenic APP processing and accumulation of β-amyloid aggregates [34].

In the current study, we show that genetic inactivation of CTSB completely abolishes accumulation of β-amyloid aggregates in the pyramidal cortical neurons of MPS IIIC *Hgsnat^P304L^* mouse model. Moreover, we demonstrate that chronic treatment with irreversible brain penetrable CTSB inhibitor E64 causes a similar effect in both MPS IIIC and sialidosis mice. The treatment also improves behavioral deficits and CNS lesions in MPS IIIC mice, suggesting that CTSB may become a novel pharmacological target for neurological LSDs.

## Results

### Cathepsin B is overexpressed and shows abnormal cytoplasmic localization in pyramidal neurons of cortical layers IV-V in the mouse models of MPS IIIC and sialidosis

To test if levels of CTSB are increased in the cortical neurons of neurological LSDs manifesting with amyloidogenesis, we analyzed levels of CTSB protein and activity in cortices of two mouse models of MPS IIIC, the HGSNAT knockout (*Hgsnat-Geo*) and knock-in (*Hgsnat^P304L^*) mouse strains, and a mouse model of sialidosis (*Neu1^τ−Ex3^*) that we have previously developed and characterized [13,35,36]. MPS IIIC patients accumulate HS, while sialidosis patients accumulate sialylated glycoproteins and oligosaccharides [37,38].

Mice were sacrificed at the age corresponding to advanced stage of CNS pathology (6 months for MPS IIIC models, 4 months for sialidosis), their brains dissected, homogenised and analyzed by immunoblotting to assess levels of CTSB protein. Our results (Figure 1A-B) indicated that in all 3 LSD models the levels of the mature 25-kDa form of CTSB were significantly increased from ∼3 (in MPS IIIC) to ∼5-fold (in sialidosis). Enzymatic CTSB activity, measured in cortex homogenates with a specific fluorogenic substrate, Z-Arg-Arg-AMC, showed a similar increase (Figure 1C).

**Figure 1.**
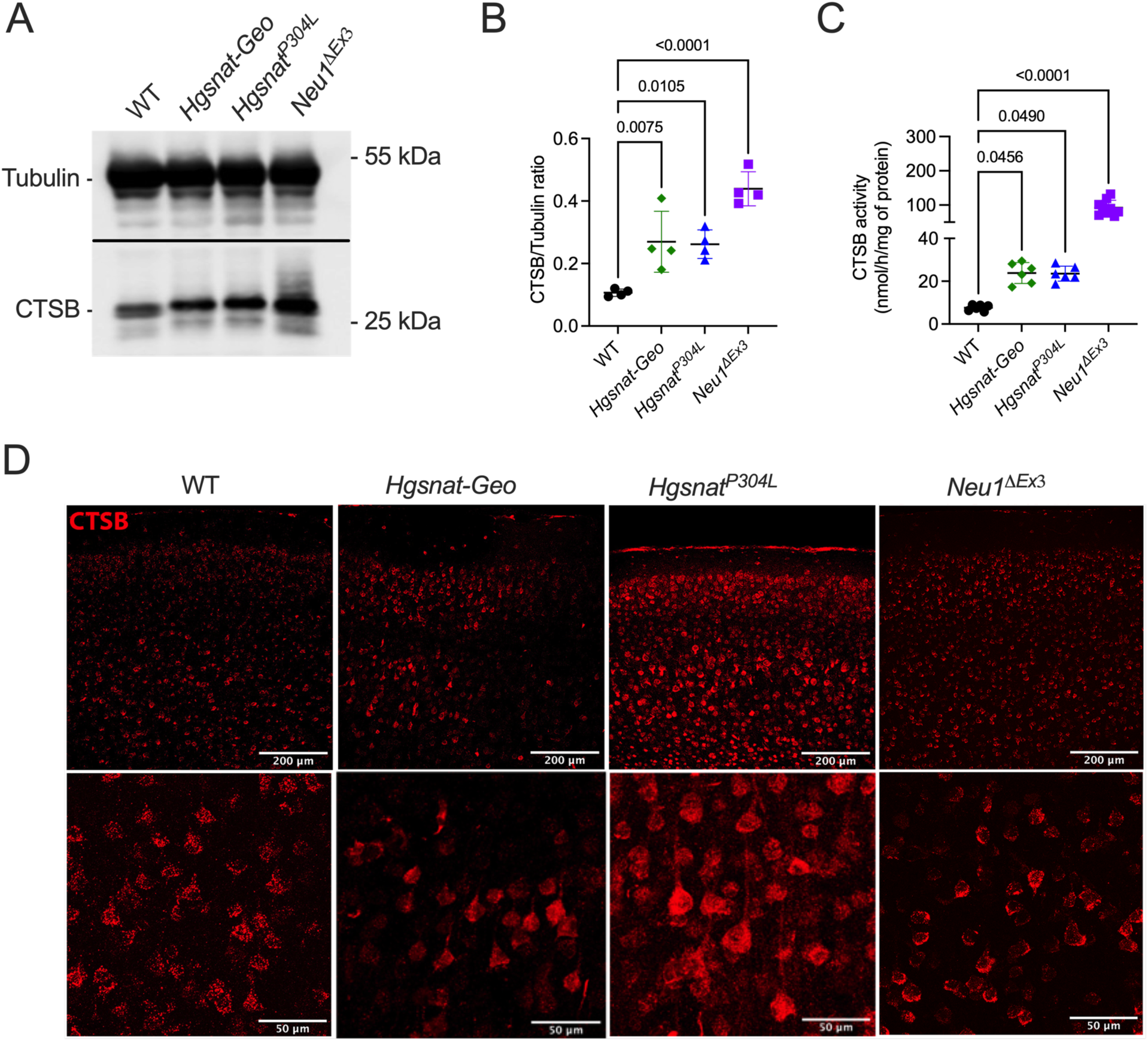
Increased CTSB levels in brain cortices of MPS IIIC and sialidosis mice. **(A)** Representative immunoblot images and **(B)** quantitative analysis of CTSB band intensities normalized by tubulin. **(C)** Enzymatic activity of CTSB in brain cortex lysates measured using the Z-Arg-Arg-AMC substrate. **(D)** Representative confocal images of brain cortex (layers IV-V) of WT, *Hgsnat-Geo*, *Hgsnat^P304L^* and *Neu1^ΔEx3^* mice showing immunolabeling for CTSB (red), analyzed at lower magnification (upper panel) and higher magnification (digital zoom of the layer V, lower panel). Individual results, means and SD from 4-9 mice per genotype are shown. P values were calculated using ANOVA with Brown-Forsythe post hoc test. Scale bars equal 200 μm (upper panels) and 20 μm (lower panels).

We further analyzed sagittal brain sections of WT and *Hgsnat-Geo* mice by immunohistochemistry to identify brain regions with the highest increase in the levels of CTSB. An increased intensity of CTSB staining in *Hgsnat-Geo* mice was observed mainly in somatosensory brain cortex, while in other areas, such as hippocampus, neurons showed levels of CTSB immunoreactivity similar to those in WT animals (supplementary Figure S1). Similar results were observed in other models which prompted us to select isocortex as the main region of interest. Further, immunofluorescent analysis of cortex tissues of all three mouse models revealed that CTSB was drastically increased in the pyramidal neurons of deep (IV-V) layers of the somatosensory cortex (Figure 1D).

CTSB levels and localization in different types of cortical cells was further studied by IHC. For this, the brain sections of WT, *Hgsnat-Geo*, *Hgsnat^P304L^* and *Neu1^τ−Ex3^* mice were labeled with antibodies against CTSB and protein markers of neurons (NeuN), astrocytes (GFAP), and activated microglia (CD11b) (supplementary Figure S2). In all studied brains of MPS IIIC and sialidosis mice, we found a drastic increase in CTSB levels in the NeuN-positive pyramidal layer IV-V neurons. The same cells also contained the highest levels of misfolded SCMAS, as well as amyloid β-positive/Thioflavin-S-positive aggregates previously identified as the robust markers of CNS pathology in neurological LSDs [13,35]. Levels of GFAP+ astrocytes in the same area were drastically increased in all models, indicative of astrocytosis, however, no co-localization was found for CTSB-positive and GFAP-positive cells suggesting that CTSB biogenesis in the astrocytes is not increased. CD11b labeling revealed multiple cells, which were also CTSB-positive and found in proximity to NeuN-positive areas, indicating that majority of activated microglia were perineuronal and that, similarly to pyramidal neurons, they also overexpressed CTSB.

Previously, we have shown in the cortical neurons of the MPS I mouse model that increased CTSB levels coincided with its cytoplasmic localization, suggesting that the enzyme was leaking from the lysosomes to the cytoplasm[34]. In the current study, diffused CTSB intracellular pattern and reduced co-localization with Lysosome Associated Membrane Protein 1 (LAMP-1) was also observed in the cortical neurons of *Hgsnat-Geo, Hgsnat^P304L^* and sialidosis mice (Figure 2A) which could be a consequence of the lysosomal membrane permeabilization. To verify this, we have stained brain sections with FITC-labeled GAL3C, the recombinant soluble lectin domain of human Galectin-3 (Figure. 2B). Previously, lysosomal recruitment of this galactose-specific lectin was shown to be associated with the lost integrity of lysosomal membranes resulting in leakage of galactose-containing proteins into the cytoplasm, and thought to be a part of the mechanism aimed on isolation of leaking lysosomes and their further elimination by autophagocytosis [39]. The pyramidal neurons in WT brains were GAL3C-negative, suggestive of preserved integrity of lysosomal membranes. In contrast, the neurons in the brains of *Hgsnat-Geo*, *Hgsnat^P304L^* and sialidosis mice contained GAL3C-positive perinuclear puncta which partially co-localized with CTSB, consistent with the leakage of the lysosomal content to the cytoplasm.

**Figure 2.**
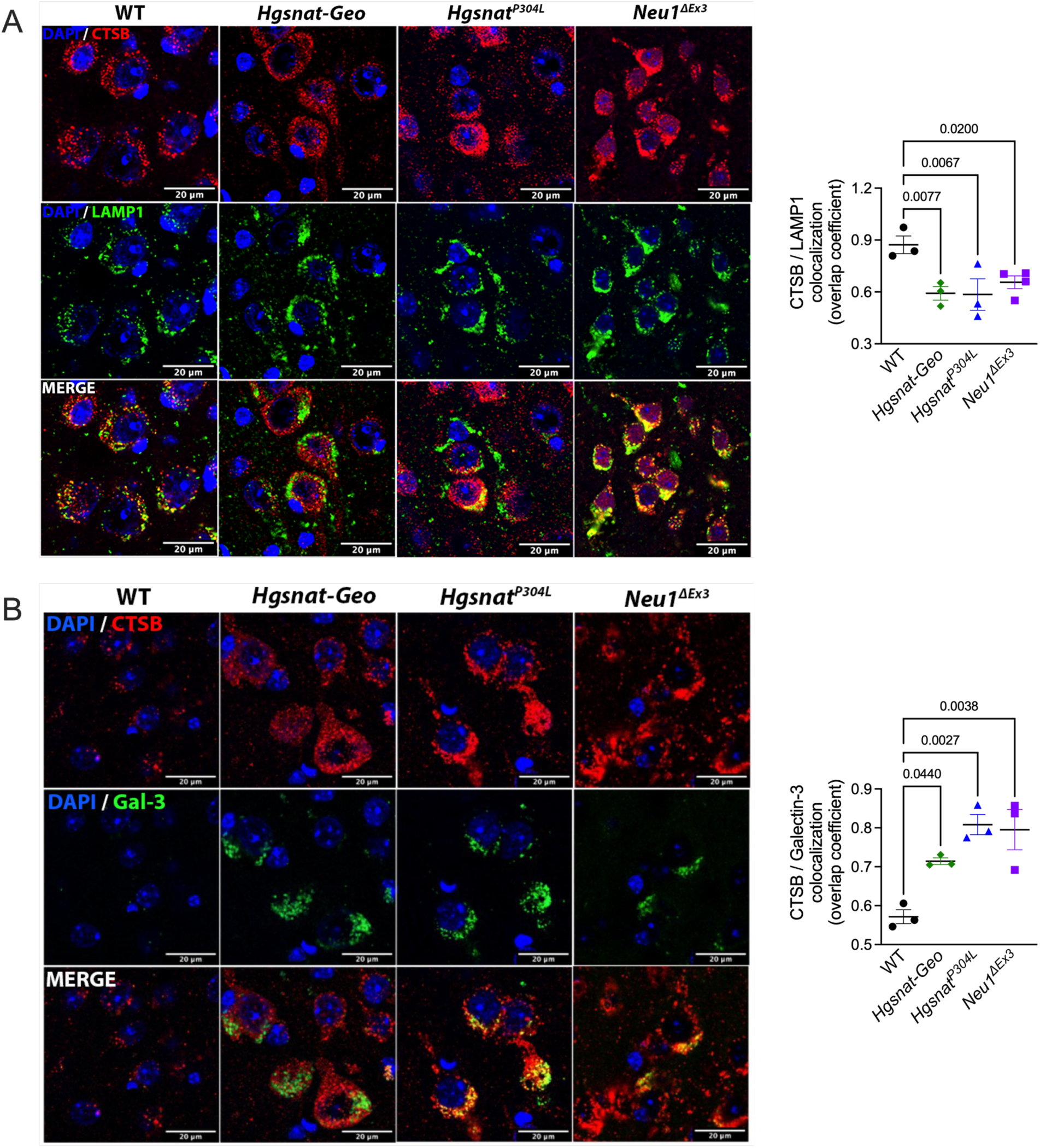
Increased levels of CTSB and its leakage to the cytoplasm in cortical neurons of MPS IIIC and sialidosis mice. Representative confocal microscopy images of brain cortex (layers IV-V) of WT, *Hgsnat-Geo*, *Hgsnat^P304L^* and *Neu1^τ−Ex3^* mice labeled for **(A)** CTSB (red) and LAMP-1 (green) or **(B)** CTSB (red) and Galectin-3 (green). DAPI (blue) was used as the nuclear counterstain. Thirty images were analyzed. Graphs show Manders’ colocalization coefficient values for CTSB and LAMP-1 or CTSB and Galectin-3 measured with ImageJ. Individual data, means and SD from 3 mice per genotype (30 images for each mouse) are shown. P values were calculated by ANOVA with Tukey post hoc test. Scale bars equal 20 μm.

The increased cytoplasmic localization of CTSB in the brain cells of *Hgsnat^P304L^* mice was further confirmed by differential centrifugation. Pooled freshly harvested brain tissues of 3 WT and 3 *Hgsnat^P304L^* 6-month-old mice were homogenized in isotonic buffer and separated to post-nuclear supernatant, organellar fraction (containing both mitochondria and lysosomes) and cytosol. The analysis of CTSB protein in the collected fractions by immunoblot (supplementary Figure S3) revealed a drastic increase of the relative CTSB content in the cytosol fraction obtained from the pooled brains of *Hgsnat^P304L^* mice (31% of the total amount detected in the post-nuclear supernatant) compared to WT mice of the same age (7.4% of the total amount) consistent with CTSB leakage from the lysosomes to the cytosol. These results also confirmed an increase in the total CTSB levels in the brains of *Hgsnat^P304L^* compared to WT mice previously revealed by enzyme activity assays and immunofluorescence microscopy (Figure S3B).

### Increased levels of CTSB are associated with presence of neuronal amyloid deposits, and amyloidogenic procession of amyloid precursor protein

In the *Hgsnat-Geo, Hgsnat^P304L^* and *Neu1^τ−Ex3^* mouse cortices, pyramidal neurons overexpressing CTSB also showed drastically increased fluorescence staining with Thioflavin-S (Figure 3A). Since after binding to beta sheet-rich peptide deposits, this dye displays enhanced fluorescence, it is commonly used for detection of misfolded proteins including amyloid aggregates in the brains of AD patients. In most neurons, Thioflavin- S staining co-localized with the areas stained with antibodies against the β-amyloid protein (AP), suggestive of amyloid accumulation in these cells (supplementary Figure S4). The levels of the full-length amyloid precursor protein (APP), measured by immunoblot in brain cortex homogenates, were similar for all groups of mice (Figure 3B). In contrast, elevated levels of 16 kDa C-terminal AP fragments (Ab 1–40 and Ab 1–42 peptides together) were found in *Hgsnat^P304L^* and *Neu1^τ−Ex3^* compared to WT mice, indicative of enhanced amyloidogenic APP processing and pro-senile neuronal phenotype. Altogether, these data suggest that the massive increase of the cytoplasmic pool of CTSB can be potentially linked to amyloidogenic processing of APP and deposition of β-amyloid aggregates in all three models.

**Figure 3.**
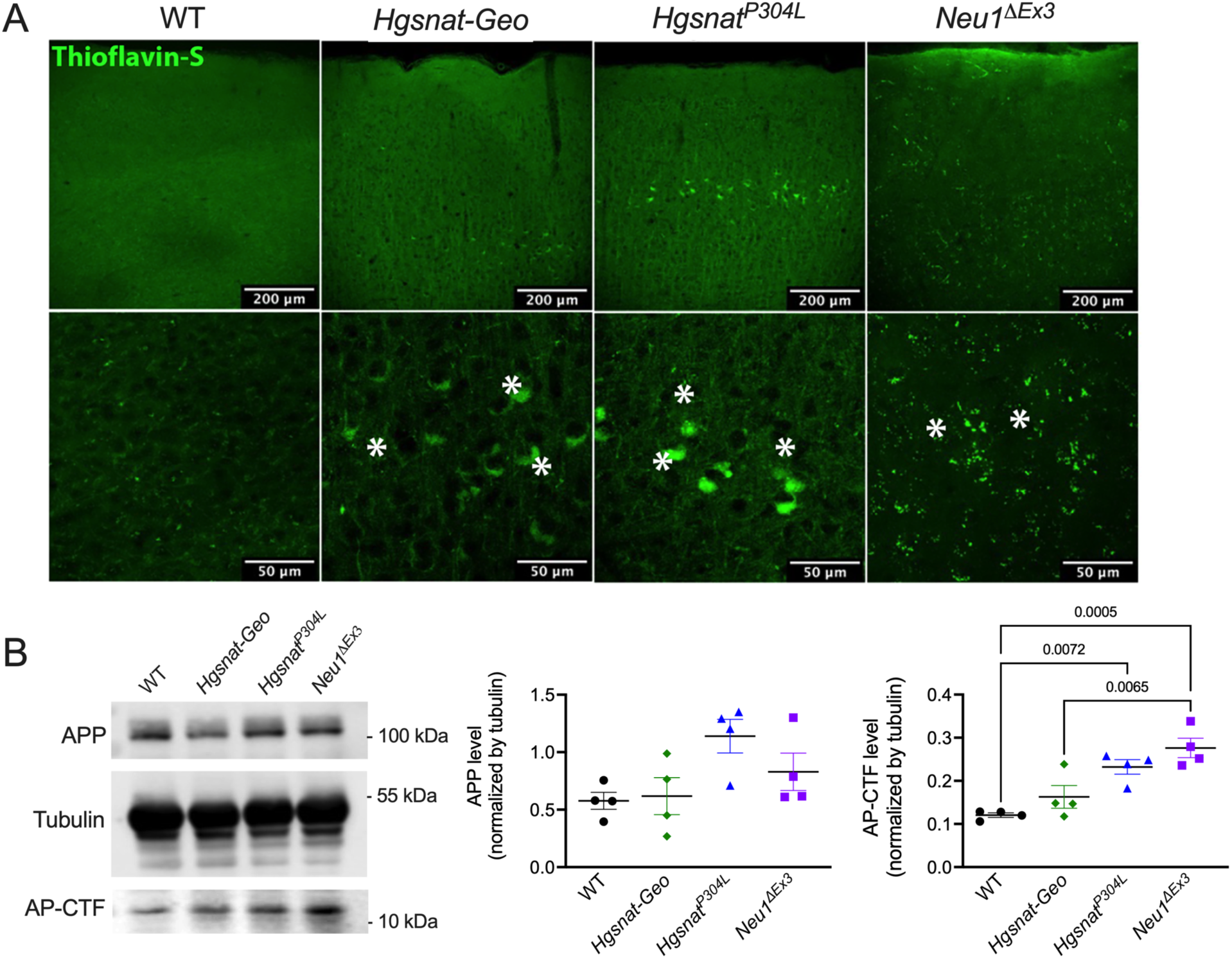
Cortices of MPS IIIC and sialidosis mice contain Thioflavin-S-positive pyramidal neurons and reveal elevated levels of 16 kDa C-terminal AP fragments. **(A)** Representative confocal microscopy images of brain cortex of MPS IIIC and sialidosis mice labeled with Thioflavin-S. Thioflavin-positive deposits in pyramidal layers IV-V neurons are marked with asterisks. Bar graphs equal 200 μm (upper panels) and 50 μm (lower panels). **(B)** Immunoblot shows unchanged levels of amyloid precursor protein (APP) but elevated levels of 16 kDa C-terminal AP fragments (AP-CTF) in cortex homogenates of *Hgsnat-Geo*, *Hgsnat^P304L^* and *Neu1^τιEx3^* mice. Graphs show individual results, means and SD. P values were calculated using ANOVA with Tukey post hoc test, n = 4 animals per genotype. Only p values <0.05 are shown.

### Levels of CTSB activity and protein are increased in cortical tissues of neurological MPS patients

To determine if the levels of CTSB and amyloid aggregates are also increased in the cortical neurons of patients affected with MPS III and other neurological MPS diseases, we analyzed frozen and PFA-fixed somatosensory cortex of post-mortem tissue, collected at autopsy and donated to the NIH NeuroBioBank. Samples from 6 MPS patients (MPS I, MPS II, MPS IIIA, MPS IIIC, and two MPS IIID) and 6 non-MPS controls, matched for age and sex, were examined (project 1071, MPS Synapse). The age, cause of death, sex, race and available clinical and neuropathological information for the patients and controls are shown in Supplementary Table S2. All MPS patients had complications from their primary disease and died prematurely (before the age of 25 years). None of the patients had received enzyme replacement therapy or hematopoietic stem cell transplantation. Enzymatic assays and immunoblotting performed on the frozen tissues confirmed that the CTSB activity and protein levels were significantly increased in cortices of MPS patients compared to controls (Figure 4A,B), suggesting that increase in the levels of this enzyme may be a common hallmark for most subtypes of MPS. In turn, staining of fixed cortical slices with Thioflavin S showed that most of patient’s tissues (4 of 5) but none of the controls were positive for the presence of misfolded protein inclusions.

**Figure 4.**
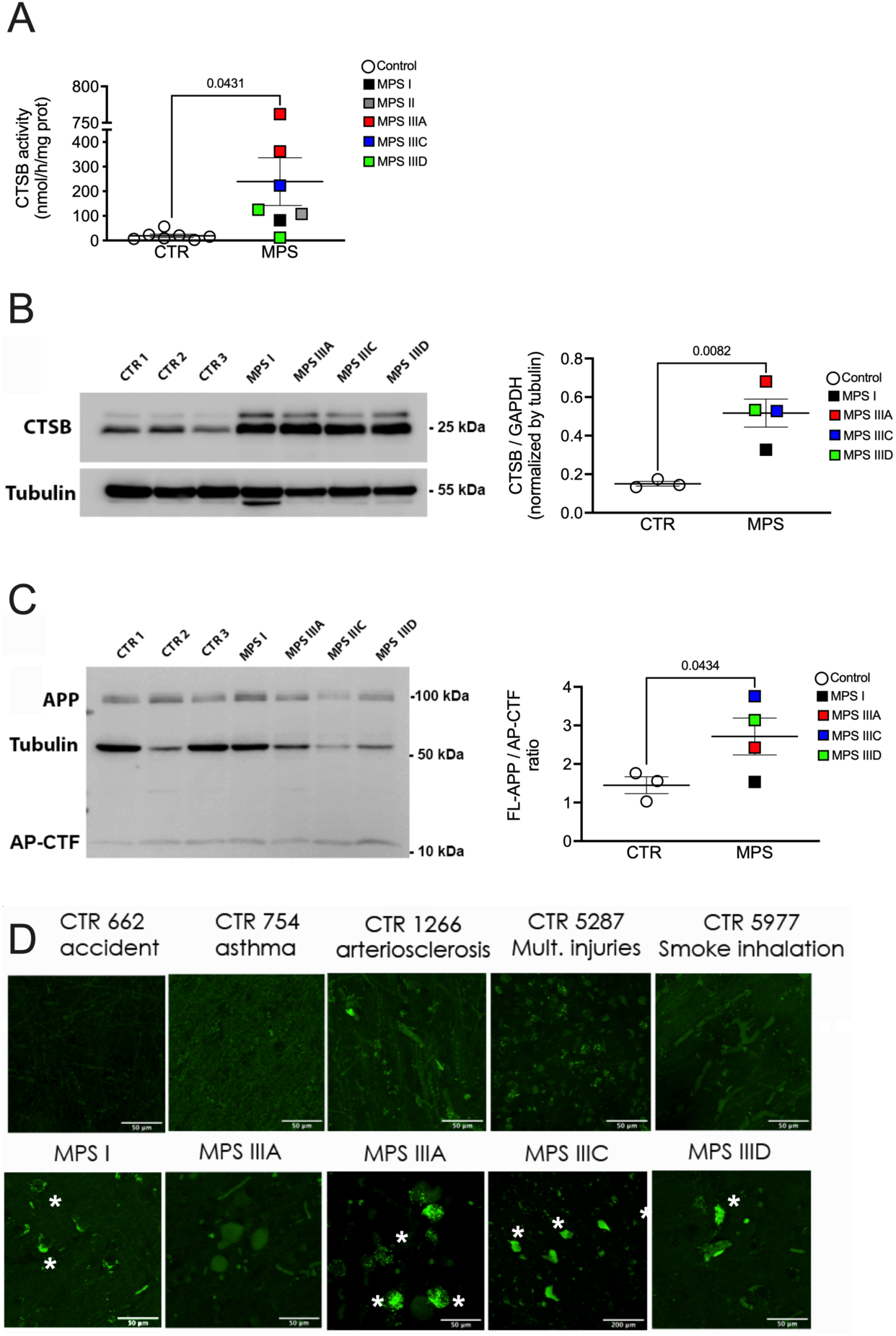
Cortices of MPS patients reveal elevated levels of CTSB activity and protein and contain Thioflavin-S-positive pyramidal neurons and elevated levels of 16 kDa C-terminal AP fragments suggestive of amyloid accumulation. **(A)** Enzymatic activity of CTSB in brain cortex lysates was measured using the Z-Arg-Arg-AMC substrate. Individual results, means and SD are shown. **(B)** Immunoblot shows elevated levels of CTSB in cortex homogenates of MPS patients compared to those without MPS (CTR). **(C)** Immunoblot shows elevated levels of 16 kDa C-terminal AP fragments (AP-CTF) in cortex homogenates of MPS patients. Graphs show individual results, means and SD. P-values were calculated by t-test. **(D)** Representative confocal microscopy images of brain cortex of MPS patients labeled with Thioflavin-S (green) show presence of misfolded protein deposits (marked with asterisks) undetectable in the cortical samples of age/sex-matching controls without MPS.

### Genetic Ctsb inactivation in the Hgsnat^P304L^ mice reduces amyloidogenesis, neuronal accumulation of G_M2_-ganglioside and neuroinflammation

To reveal a causative relation between the increase in the CTSB levels and accumulation of amyloid aggregates in the cortical neurons, we studied *Hgsnat^P304L^* mice with genetic depletion of the *Ctsb* gene (*Hgsnat^P304L^*/*Ctsb^-/-^*). The strain was generated by breeding the *Hgsnat^P304L^* strain with previously described *Ctsb* KO mice (Jax Strain [B6;129- Ctsbtm1Jde/J]). Analysis of the brain tissues of *Ctsb^-/-^* and *Hgsnat^P304L^*/*Ctsb^-/-^*mice demonstrated an absence of CTSB enzymatic activity (Figure 5A) or immunoreactive material (Figure 5B). *Ctsb^-/-^* mice do not show increased levels and size of LAMP-2- positive puncta in the neurons, a marker of lysosomal storage (Figure 5C), and do not display biomarkers of CNS pathology common for neurological LSDs, including astromicrogliosis (Figure 5D). Analysis of CNS pathology in 6-month-old *Hgsnat^P304L^*/*Ctsb^-/-^* mice revealed drastically reduced levels of Thioflavin-S/β-amyloid double-positive aggregates (Figure 5E) in cortical pyramidal neurons compared to *Hgsnat^P304L^* mice, demonstrating that depletion of CTSB blocked amyloidogenesis. Besides, the cortical neurons of *Hgsnat^P304L^*/*Ctsb^-/-^* mice showed reduced levels of misfolded SCMAS (Figure 5F) and G_M2_ ganglioside (Figure 5G) and did not contain LC3-positive and P62-positive puncta suggesting that CTSB depletion also rescued an autophagy block (Figure 5H and Figure 5K). On the other hand, the level of astromicrogliosis in the CTSB-deficient *Hgsnat^P304L^*/*Ctsb^-/-^* mice was similar to that in the *Hgsnat^P304L^* strain (Figure 5D).

**Figure 5.**
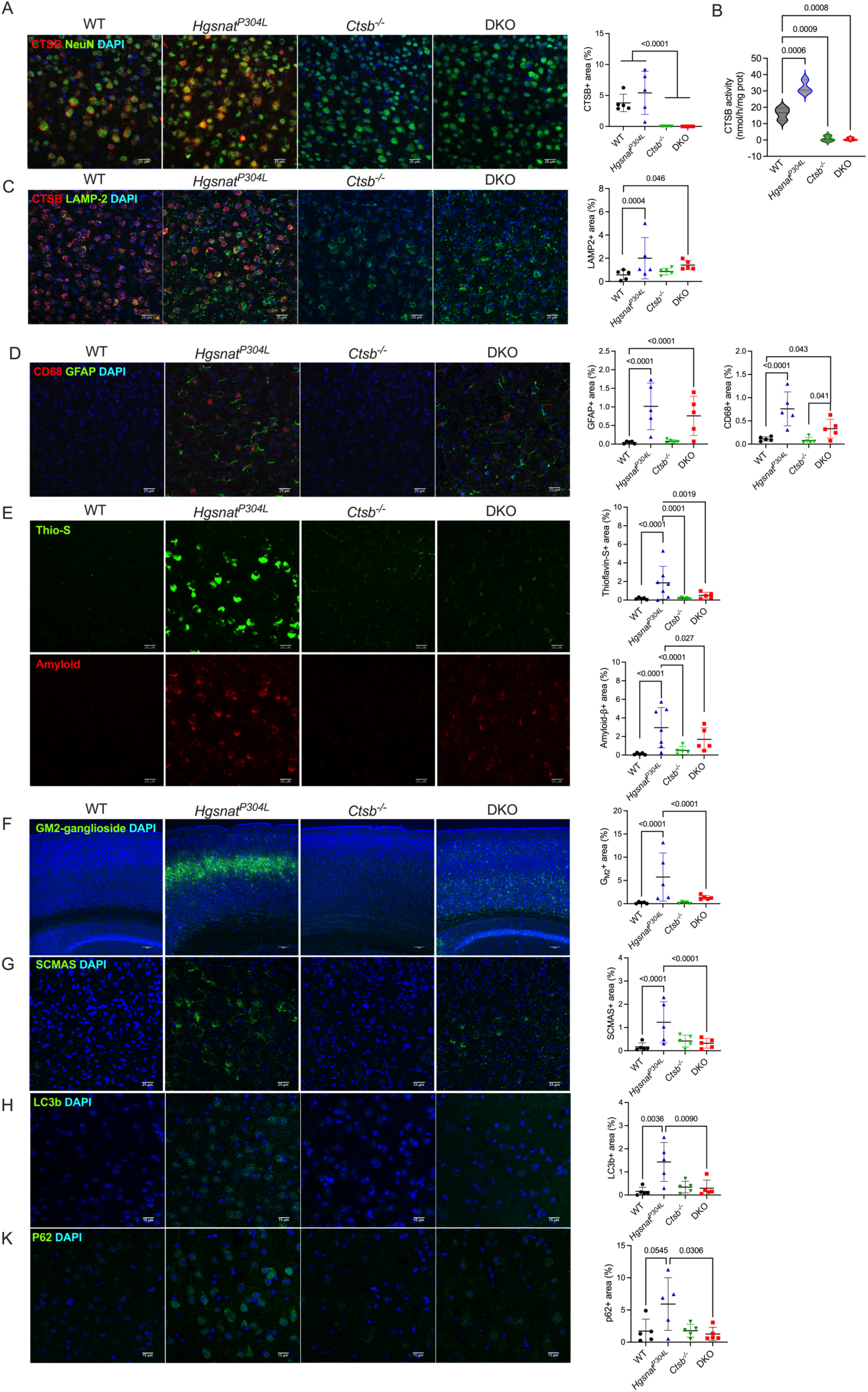
Genetic inactivation of CTSB reduces levels of Thioflavin S-positive/β-amyloid-positive aggregates and restores autophagic flux in cortical neurons of MPS IIIC mice. **(A-B)** CTSB immunoreacting material **(A)** and enzymatic activity **(B)** are not detected in cortical tissues of *Ctsb^-/-^* and *Hgsnat^P304L^*/*Ctsb^-/-^* (DKO) mice. (C) *Hgsnat^P304L^* and *Hgsnat^P304L^*/*Ctsb^-/-^* mice show increased levels and size of LAMP-2-positive puncta in the neurons. **(D)** 6-month-old *Hgsnat^P304L^* and *Hgsnat^P304L^*/*Ctsb^-/-^* mice show similar levels of GFAP-positive (green) and CD68-positive (red) cells. **(E)** Storage of β-amyloid (Amyloid, red) and Thioflavin-S-positive misfolded proteins (Thio-S, green) is elevated in cortical neurons of 6-month-old *Hgsnat^P304L^* mice but not in double knockout *Hgsnat^P304L^*/*Ctsb^-/-^* mice. **(F-K)** Genetic ablation of *Ctsb* reduces accumulation of GM2-ganglioside **(F)**, SCMAS **(G)**, LC3-positive **(H)** and p62-positive **(K)** puncta in cortical neurons of *Hgsnat^P304L^* mice. Bar graphs equal 100 μm in **(F)**, 15 μm in **(H,K)** and 25 μm in all other panels. Panels show representative confocal microscopy images of brain cortices (layers IV-V) of WT, *Ctsb^-/^,^-^ Hgsnat^P304L^*, and *Hgsnat^P304L^*/*Ctsb^-/-^* mice. Graphs show quantification of immunofluorescence with ImageJ software. Individual data, means and SD for 5-7 mice per genotype (3 sections for each mouse) are shown. Statistical analysis was performed by nested ANOVA with Tukey post hoc test.

### Chronic treatment of Hgsnat^P304L^ mice with a pharmacological CTSB inhibitor E64 rescues behavioural abnormalities and reduces amyloid deposits in cortical pyramidal neurons

Taking advantage of the fact that the brain permeable irreversible CTSB inhibitor E64 (also known as Aloxistatin) is available and does not show toxicity in the doses of 1- 5 mg/kg BW, sufficient to inhibit the enzyme in the mouse brain tissues [40–43], we further tested if pharmacological CTSB inhibition matches the results of its genetic inactivation in *Hgsnat^P304L^* mice. As a delivery route, we have chosen an intranasal administration, which is non-invasive and showed efficacy in delivery of small (<1000 Da) molecules to the brain parenchyma [44]. Mice were treated with daily E64 doses of 1 mg/kg BW between the ages of 5 and 6 months, i.e., directly preceding the age when accumulation of amyloidogenic deposits in cortical neurons becomes evident [35]. Mice were then analysed by an Open Field (OF) test, instrumental to detect hyperactivity and reduced anxiety characteristic of MPS IIIC mice, and sacrificed to analyse CNS pathology. Similar analyses were also conducted for untreated *Hgsnat^P304L^* and *Hgsnat^P304L^*/*Ctsb^-/-^*mice as well as for untreated and treated WT mice of the similar age and sex. As expected, untreated 6-month-old *Hgsnat^P304L^* mice showed an increase in the total travel distance (hyperactivity, Figure 6A) and in the amount of time spent in the central area (reduced anxiety, Figure 6B-F) compared to WT mice. In contrast, *Hgsnat^P304L^* mice treated with E64 demonstrated a normal behaviour in the OF test similar to that of the control mice. WT mice treated with E64 showed behaviour undistinguishable from that of untreated WT mice. Somewhat unexpectedly, both *Hgsnat^P304L^*/*Ctsb^-/-^* and *Ctsb^-/-^* mice, similarly to untreated *Hgsnat^P304L^* mice, showed an increase in total distance traveled and time spent/distance traveled in the central area compared to WT mice consistent with hyperactivity and reduced anxiety.

**Figure 6.**
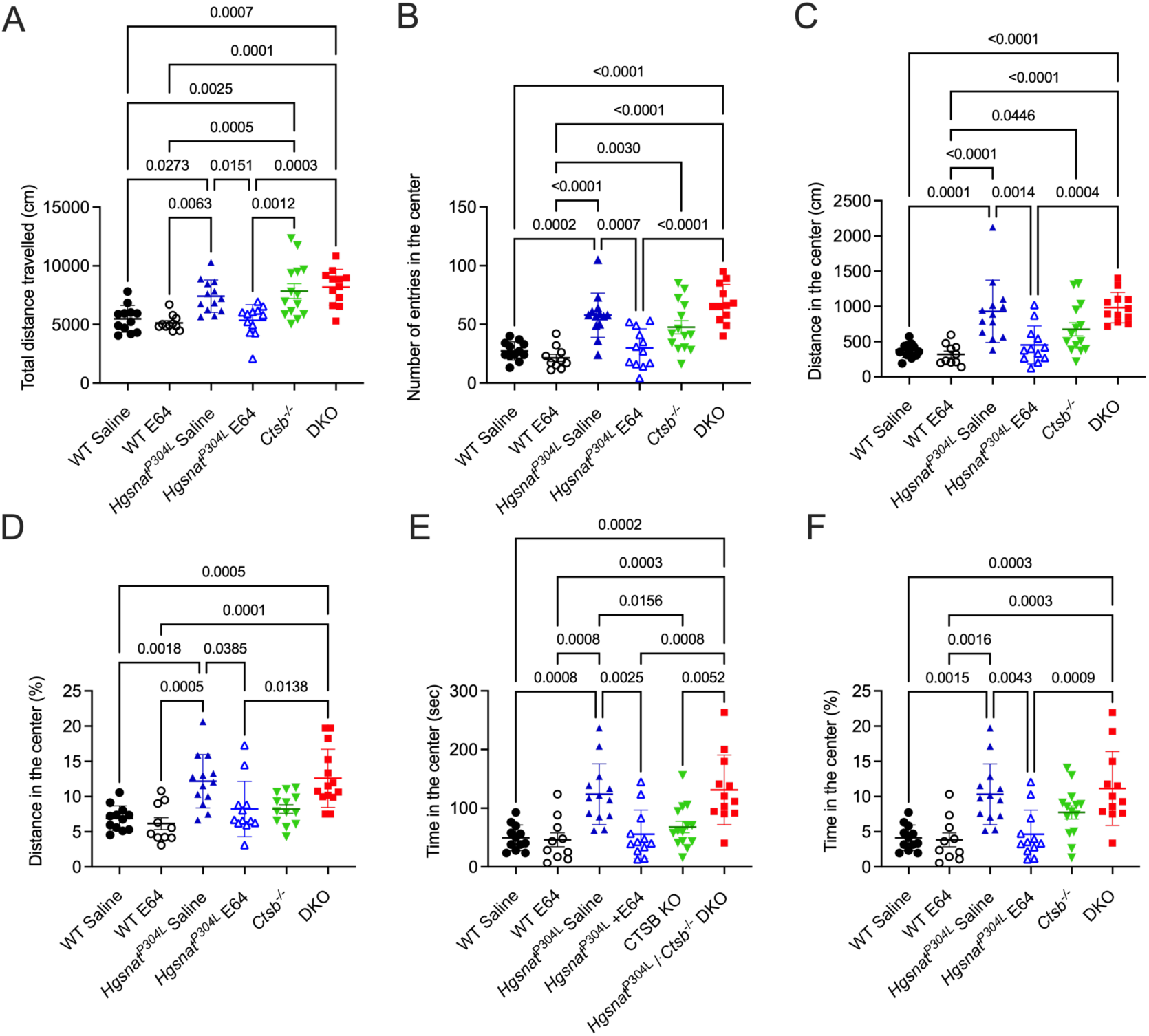
Thirty-day intranasal treatment with E64 rescues behavioural deficits in MPS IIIC mice. *Hgsnat^P304L^* mice treated intranasally with E64 for 30 days at 1 mg/kg BW/day show significant reduction of hyperactivity **(A)** and reduced anxiety **(B-F)** in the Open Field test at the age of 6 months compared with untreated *Hgsnat^P304L^* mice. Double-deficient *Hgsnat^P304L^*/*Ctsb^-/-^* mice show behavior similar to that of untreated *Hgsnat^P304L^* mice. Individual results, means and SD from experiments performed with 10-14 mice per genotype, per treatment are shown. P values were calculated by ANOVA with Tukey post hoc test.

Analysis of the brain tissues by IHC conducted in 7-month-old mice revealed that, like the *Hgsnat^P304L^*/*Ctsb^-/-^* mice, E64-treated *Hgsnat^P304L^* mice did not contain Thioflavin-S-positive and APP-positive depositions in cortical neurons (Figure 7A), thus demonstrating that the drug effectively prevented accumulation of amyloid aggregates. E64-treated *Hgsnat^P304L^* mice also did not show SCMAS storage (Figure 7B) as well as LC3- positive and P62-positive puncta in cortical pyramidal neurons (Figure 7C) suggesting that the drug restored normal autophagy levels.

**Figure 7.**
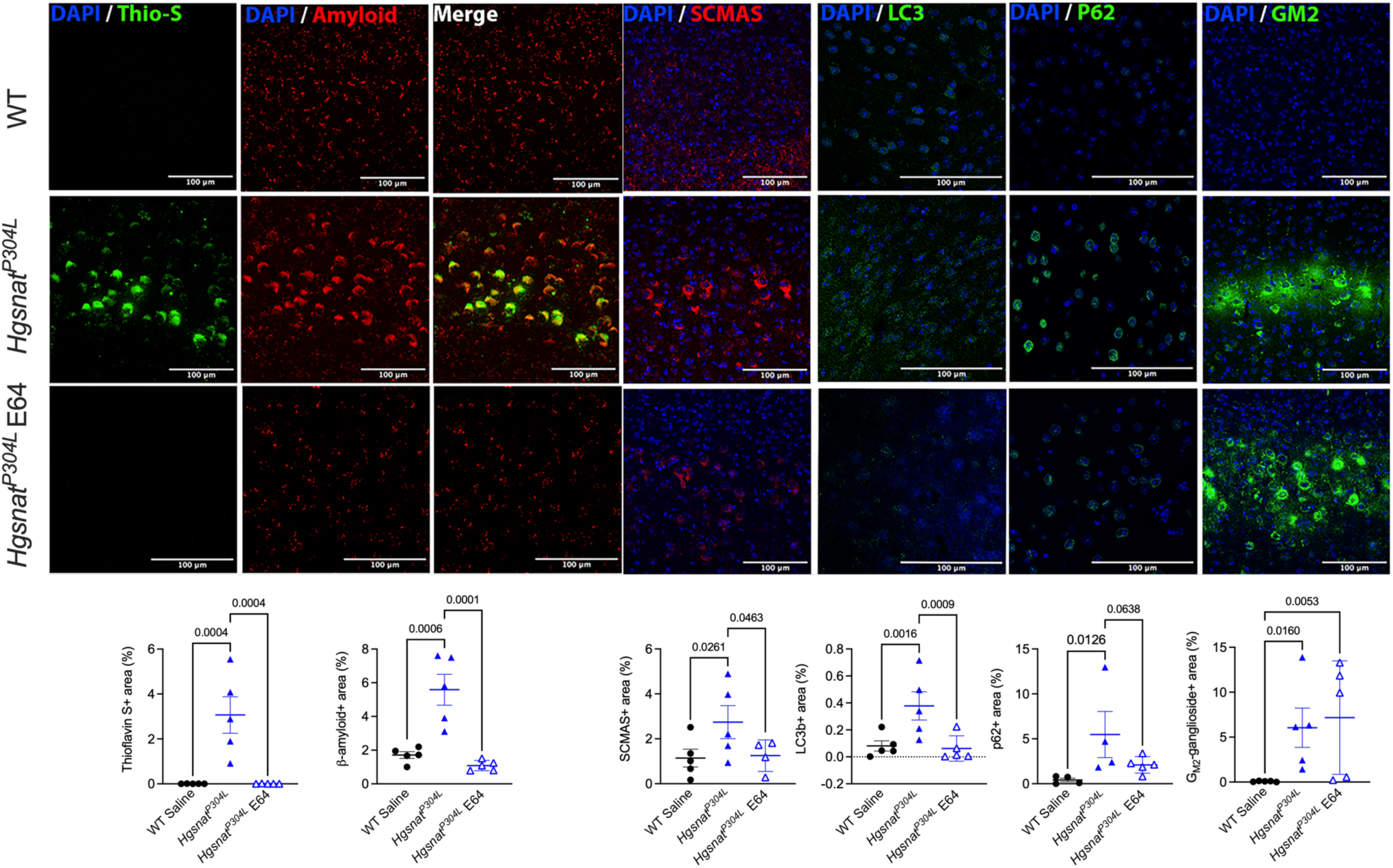
Thirty-day intranasal treatment with irreversible CTSB inhibitor E64 reduces levels of Thioflavin S-positive/β-amyloid-positive aggregates and restores autophagic flux in cortical neurons of MPS IIIC mice. **(A)** Accumulation of β-amyloid and misfolded proteins is not detected in cortical neurons of 6-month-old *Hgsnat^P304L^* mice treated for 30 days with daily intranasal doses of E64 at 1 mg/kg BW. **(B-D)** E64 treatment reduces accumulation of SCMAS **(B)**, LC3-positive **(C)** and p62-positive **(D)** puncta but not secondary storage of GM2-ganglioside **(E)** in cortical neurons of *Hgsnat^P304L^* mice. Panels show representative confocal microscopy images of brain cortices (layers IV-V) of WT, untreated *Hgsnat^P304L^*, and E64-treated *Hgsnat^P304L^* mice labeled for β-amyloid (**A**, red), Thioflavin-S (**A**, green), SCMAS (**B**, red) LC3 (**C**, green), p62 (**D**, green), and GM2-ganglioside (**E**, green). Graphs show quantification of immunofluorescence with ImageJ software. Individual data, means and SD are shown. Statistical analysis was performed by ANOVA with Tukey post hoc test; n = 5 mice per genotype.

At the same time, like in CTSB-deficient *Hgsnat^P304L^*/*Ctsb^-/-^* mice, the level of astromicrogliosis in the *Hgsnat^P304L^* mice treated with E64 was similar to that in the untreated *Hgsnat^P304L^* mice (supplementary Figure S5). This was consistent with increased expression levels of pro-inflammatory cytokines MIP1α, and TNFα in their brain tissues (supplementary Figure S6). On the other hand, expression of IL-1β was normalised in the brain of treated mice.

### Chronic treatment of sialidosis mice with E64 reduces amyloid deposits in cortical layer 4-5 pyramidal neurons

We further tested whether E64 treatment would also reduce amyloidogenesis in the cortices of sialidosis mice. Since these mice show more aggressive disease progression causing severe debilitation and death by the age of 18-24 weeks [36], we treated them starting from weaning (∼6 weeks) and until the age of 16 weeks, when the mice were sacrificed and their fixed brain tissue cryopreserved to study accumulation of amyloid deposits in the cortical neurons. This analysis revealed that the β-amyloid/Thioflavin-S-positive aggregates, clearly present in the pyramidal cortical layer IV-V neurons of untreated *Neu1^ΔEx3^* mice, were absent in the brains of both WT and E64-treated *Neu1^ΔEx3^* mice (Figure 8A). Similarly to E64-treated MPS IIIC *Hgsnat^P304L^* mice and MPS IIIC mice with CTSB deficiency (*Hgsnat^P304L^*/*Ctsb^-/-^*), E64-treated sialidosis mice also showed a significantly reduced levels of SCMAS aggregates in the same type of neurons, not statistically different from their WT counterparts (Figure 8B). Together, these data are suggestive of CTSB involvement in amyloidogenesis in different types of neurological LSDs.

**Figure 8.**
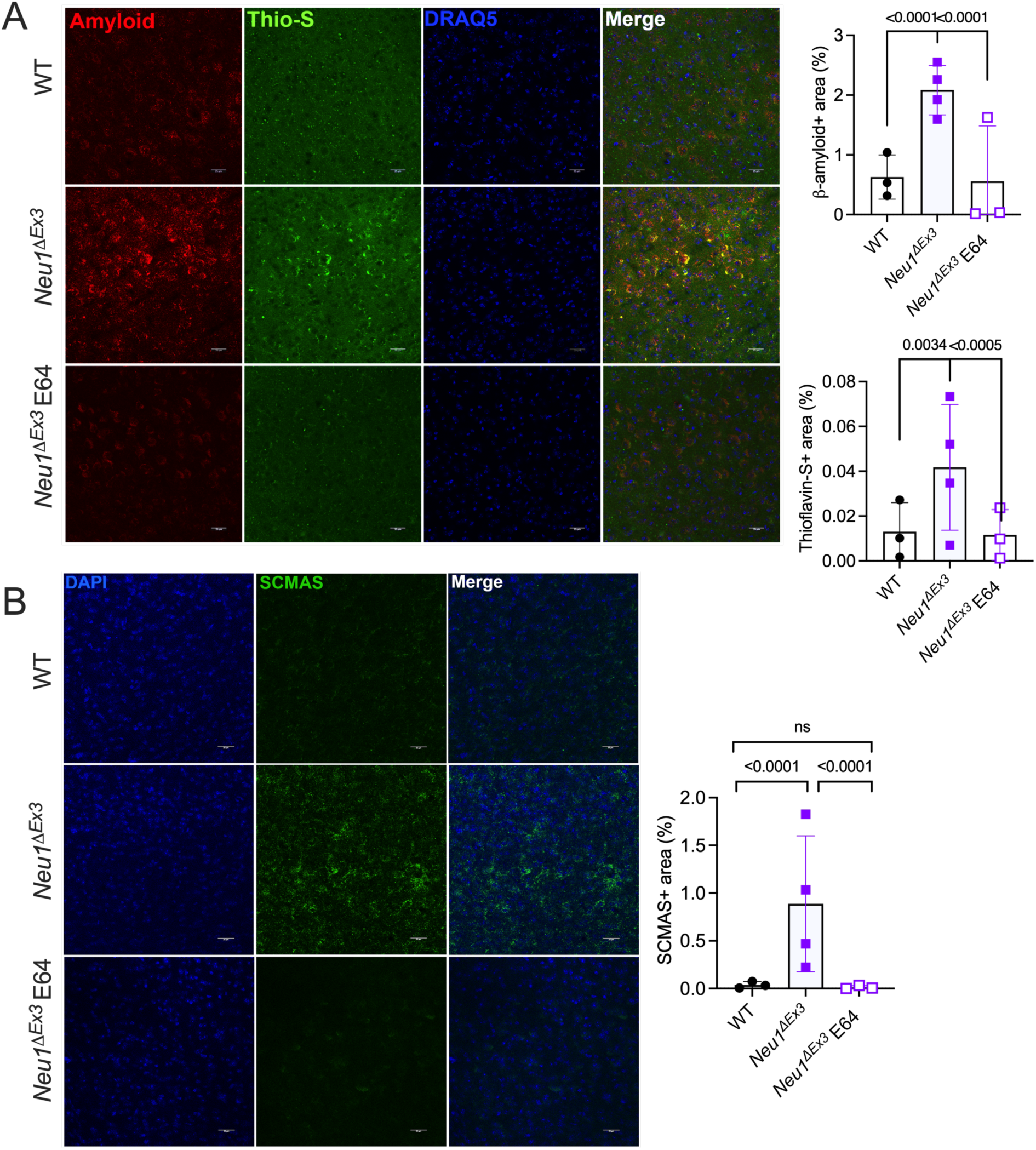
Thirty-day intranasal treatment with irreversible CTSB inhibitor E64 reduces levels of Thioflavin S-positive/β-amyloid-positive aggregates and misfolded SCMAS deposits in cortical neurons of sialidosis mice. **(A)** Storage of amyloidogenic peptides and misfolded proteins reduced in cortical neurons of 4-month-old *Neu1^ΔEx3^* mice treated for 60 days with daily intranasal doses of E64 at 1 mg/kg BW. **(B)** E64 treatment also reduces accumulation of SCMAS in cortical neurons of *Neu1^ΔEx3^* mice. Panels show representative confocal microscopy images of brain cortices (layers IV-V) of WT, *Neu1^ΔEx3^* , and E64-treated *Neu1^ΔEx3^* mice labeled for β-amyloid (**A**, red), Thioflavin-S (**A**, green), and SCMAS (**B**, green). Nuclei were counterstained with Draq5 (**A**, blue) and DAPI (**B**, blue). Graphs show quantification of immunofluorescence with ImageJ software. Individual data, means and SD are shown. Statistical analysis was performed by nested ANOVA with Tukey post hoc test; n=3-4 mice per genotype. Four brain sections were analysed for each mouse.

## Discussion

Despite many years of extensive research, no specific treatments have been clinically approved for MPS III and sialidosis. Moreover, as we currently know, gene therapy alone cannot reverse CNS pathology in the symptomatic patients emphasizing the need for development of additional/alternative strategies and, in particular, those using small molecule drugs [45,46]. In the current work, we examined whether inhibition of CTSB, which we previously proposed to be responsible for the increased amyloidogenesis in neurological MPS diseases [34], could ameliorate disease progression in MPS IIIC and sialidosis mouse models. Together, our data provide compelling evidence that secondary induction of CTSB in the pyramidal neurons contribute to accumulation of amyloid deposits and misfolded proteins in neurological lysosomal diseases, MPS IIIC and sialidosis. We also demonstrate that by inhibiting CTSB activity we can partially improve behavioral deficits and CNS pathology in MPS IIIC mice.

To establish a causative relation between the increased levels of CTSB and increased amyloidogenesis, we generated a knock-in MPS IIIC mouse model with inactivated *Ctsb* gene (*Hgsnat^P304L^*/ *Ctsb^-/-^* strain). According to published studies, *Ctsb^-/-^* mice do not reveal severe neurological phenotype or reduced lifespan suggesting that CTSB function in the lysosome is somewhat redundant, and its deficiency can be partially compensated by other proteases [47,48]. We also treated MPS IIIC mice with an irreversible inhibitor of CTSB and other cysteine proteases, E64, at the dose and frequency previously reported to be sufficient to cause a sustainable inhibition of CTSB in the brain [40–43]. In both cases, we have observed a drastic reduction of β-amyloid- positive/Thioflavin S-positive cytoplasmic aggregates in pyramidal neurons of deep cortical layers IV and V, the cells showing the highest levels of both amyloid deposits and CTSB immunolabeling in untreated *Hgsnat^P304L^* mice and in *Hgsnat-Geo* mice, a knockout model of MPS IIIC. Similarly to our previous findings in *Idua^-/-^* MPS I mice [34], CTSB shows a diffused labeling pattern in pyramidal cortical neurons of both sialidosis and MPS IIIC mice which, together with the presence of GAL3C-positive puncta in these neurons, and increased CTSB content in the cytosol fraction from *Hgsnat^P304L^* mouse brain tissues is suggestive of CTSB leakage from the lysosome to the cytoplasm. In contrast to other lysosomal cysteine proteases, CTSB remains stable at neutral pH corresponding to that of the cytoplasm (reviewed in ref. [49]). Moreover, a recent report demonstrates that CTSB, which mainly shows carboxypeptidase and dipeptidyl carboxypeptidase activities at the acidic pH of the lysosome, reveals endopeptidase activity at the neutral pH [50], which presumably allows the enzyme to conduct amylogenic processing of APP.

Notably, in the neurons of both E64-treated and *Ctsb*-depleted MPS IIIC mice, we also detected a drastic reduction of other secondary storage product, misfolded SCMAS protein, which accumulation have been previously associated with an autophagy block. In contrast to neurons of untreated *Hgsnat^P304L^* mice, the cells in both E64-treated and CTSB KO *Hgsnat^P304L^* mice did not display LC3-positive and P62-positive puncta suggesting that CTSB depletion indeed rescued an autophagy flux. This is also consistent with the results of a recent study showing that CTSB regulates autophagy through cleavage of the lysosomal calcium channel MCOLN1/TRPML1 [33].

At the same time, the effect of E64 treatment on behavioural abnormalities in *Hgsnat^P304L^* mice was different from that of genetic *Ctsb* depletion. E64-treated mice demonstrated a rescue of hyperactivity and reduced anxiety characteristic for Sanfilippo mouse models [12,13,51]. In contrast, the *Hgsnat^P304L^/Ctsb^-/-^* mice showed levels of hyperactivity and reduced anxiety similar to those of untreated *Hgsnat^P304L^* mice. E64 treatment did not cause any changes in behaviour of control WT mice outruling the possibility that reduction of hyperactivity in the *Hgsnat^P304L^* mice was caused by a sedative effect of the drug itself. Notably, while we do not see signs of CNS pathology, characteristic for neurological LSDs, such as astromicrogliosis related to neuroinflammation, or neuronal accumulation of gangliosides and ceroid materials in *Ctsb^-/-^* mice, their levels of hyperactivity and reduced anxiety are comparable to those of *Hgsnat^P304L^* mice. Therefore, in the current experimental settings, it was impossible for us to conclude whether the above behaviour deficits in *Hgsnat^P304L^/Ctsb^-/-^* mice originated from CTSB genetic deficiency, HGSNAT deficiency or both. Previously, CTSB genetic depletion has been shown to ameliorate behavioral deficits, neuropathology, and biomarkers in several mouse models of neurological diseases including traumatic brain injury, ischemia, epilepsy, multiple sclerosis, opioid tolerance, and inflammatory pain (reviewed in ref. [47]). At the same time, a recent study demonstrated that in iPSC-derived dopaminergic neurons CTSB depletion reduced lysosomal glucocerebrosidase activity and impaired the degradation of pre-formed alpha-synuclein fibrils associated with pathogenesis in Parkinson disease [52]. Thus, the effect of CTSB reduction can be both positive and negative depending on the target cells and degree of depletion.

Our findings that treatment of MPS IIIC mice with the CTSB inhibitor E64 effectively prevents accumulation of amyloid materials and rescues behavioral abnormalities positions this drug as a promising candidate for clinical translation in MPS IIIC and perhaps other neurological lysosomal diseases. Although the mechanism of the drug action needs further elucidation, the fact that it reduces accumulation of secondary storage materials suggests that, besides blocking amyloidogenic cleavage of APP, E64 also induces autophagy. The drug also reduced expression of proinflammatory cytokine IL-1β consistent with its role in the inflammasome activation (reviewed in ref. [53]). At the same time, E64 treatment does not reduce astromicrogliosis and expression levels of two other proinflammatory cytokines MIP1α, and TNFα. Notably, a recent clinical trial demonstrated the improved neurobehavioral and functional outcomes in symptomatic MPS III patients undergoing an anti-inflammatory therapy [54]. We hypothesize, therefore, that for an effective treatment of progressive neuropathology, E64 needs to be used in combination with a therapy reducing neuroimmune response, such as, for example, hematopoietic stem progenitor cell (HSPC) transplantation or HSPC gene therapy. Experiments aimed on testing this hypothesis are now in progress in our laboratory. Previously E64 has shown efficacy in rescuing memory loss and reducing amyloid plaque formation in the AD mouse model [43], demonstrated oral bioavailability and has been approved by FDA for clinical trials. Pharmacological CTSB inhibitors also ameliorated neurodegeneration after traumatic brain injury [55], retinopathy and optic neuritis in experimental autoimmune encephalomyelitis [56], or HIV-1 induced neuronal damage [57]. Besides, potential indications for CTSB inhibiting drugs include other neurological and psychiatric diseases associated with increased CTSB levels such as amyotrophic lateral sclerosis, bipolar disorder, attention deficit hyperactivity disorder and autism spectrum (reviewed in ref. [47,53]).

Notably, multiple clinical trials involving enzyme replacement or gene therapy for MPS patients, both completed and ongoing, have yet to show efficacy for symptomatic patients suggesting that correction of the enzymatic defect by itself cannot reverse or even halt progression of neuronal pathology. Our current results showing that E64 administered by non-invasive intranasal route blocked amyloidogenesis in both symptomatic MPS IIIC and sialidosis mice justify further studies analysing efficacy of this compound for treatment of neurological LSDs.

## Materials and Methods

### Study approval

All animal experiments were approved by and performed in compliance with the Comité Institutionnel des Bonnes Pratiques Animales en Recherche (CIBPAR; approval number 2022-4353, approval date November 2023). Ethical approval for research involving human tissues was given by the CHU Ste-Justine Research Ethics Board (Comité d’éthique de la recherche FWA00021692, approval number 2020-2365). The National Institute of Health’s NeuroBioBank provided the cerebral tissues, frozen or fixed with paraformaldehyde, from MPS patients as well as age, ethnicity and sex-matched controls (project 1071, MPS Synapse), along with clinical descriptions and the results of a neuropathological examination.

### Animals

The constitutive *Neu1* KO sialidosis mouse model *Neu1^ΔEx3^*, the constitutive knockout (KO) MPS IIIC mouse model *Hgsnat-Geo* and knock-in MPS IIIC mouse model *Hgsnat^P304L^*, expressing the HGSNAT enzyme with human missense mutation P304L, both on a C57BL/6J genetic background, have been previously described [13,51,58]. The strain of *Hgsnat^P304L^*/*Ctsb^-/-^* mice was generated by breeding the *Hgsnat^P304L^* strain with previously described *Ctsb* KO mice obtained from the Jackson Laboratory (Jax Strain [B6;129- Ctsbtm1Jde/J]). Heterozygous mice were interbred, and litters were genotyped by PCR using genomic DNA extracted from clipped tail tip, as described. Mice homozygous for the mutant allele(s) were compared to appropriate age- and sex-matched wild-type (WT) control littermates. All mice were housed under 12 h/12 h light - dark cycles with ad libitum access to a normal rodent chow and water.

Equal cohorts of male and female mice were studied separately for each experiment, and statistical methods were used to test whether the progression of the disease, levels of biomarkers or response to therapy were different for male and female animals. Since differences between sexes were not detected, the data for male and female mice were pooled together.

### Immunohistochemistry

Mouse brains were collected from animals, perfused with 4% PFA in PBS and post-fixed in 4% PFA in PBS overnight. Brains were incubated in 30% sucrose for 2 days at 4°C, embedded in Tissue-Tek® OCT Compound, cryopreserved, cut in 40 µm-thick sections and stored in cryopreservation buffer (0.05 M sodium phosphate buffer, pH 7.4, 15% sucrose, 40% ethylene glycol) at −20°C pending immunohistochemistry. Mouse brain sections were washed 3 times with PBS and permeabilized/blocked by incubating in 5% bovine serum albumin (BSA), 0.3% Triton X-100 in PBS for 1 h at room temperature. Incubation with primary antibodies, diluted in 1% BSA, 0.3% Triton X-100 in PBS, was performed overnight at 4°C. The antibodies and their working concentrations are shown in the supplementary Table S1.

To conduct Thioflavin-S staining, brain sections stained with Draq5, primary antibodies against β-amyloid and Alexa Fluor-labeled secondary antibodies were washed 3 times with PBS and incubated in a 0.05% Thioflavin-S (Sigma, T1892) solution in 50% ethanol/water for 10 min protected from light. Then the sections were washed 2 times with 50% ethanol followed by two washes with ddH_2_O.

The slides were mounted with Prolong Gold Antifade mounting reagent (Invitrogen, P36934) and analyzed using Leica DM 5500 Q upright confocal microscope (10x, 40x, and 63x oil objective, N.A. 1.4). Autofluorescent ceroid materials in brain cortices were analysed using mounted unstained tissue sections and confocal settings similar to those for Green Fluorescent Protein. Images were processed and quantified using ImageJ 1.50i software (National Institutes of Health, Bethesda, MD, USA) in a blinded fashion. Panels were assembled with Adobe Photoshop.

### Enzymatic assays and western blots

CTSB activity was measured using the fluorogenic Z-Arg-Arg-AMC substrate (Enzo Life Sciences, USA) as described by Viana et al. [34]. Levels of CTSB protein, total amyloid precursor proteins (APP) and C-terminal AP fragments (AP-CTF, Ab 1–40 and Ab 1–42 peptides together) were analysed by immunoblots as previously described [34].

### Real-time qPCR

RNA was isolated from snap-frozen brain tissues using the TRIzol reagent (Invitrogen) and reverse-transcribed using the Quantitect Reverse Transcription Kit (Qiagen # 205311) according to the manufacturer’s protocol. qPCR was performed using a LightCycler® 96 Instrument (Roche) and SsoFast™ EvaGreen® Supermix with Low ROX (Bio RAD #1725211) according to the manufacturer’s protocol. The primers are shown in the supplementary Table S2. The relative levels of mRNA were calculated using the 2-ΔΔCt algorithm with RLP32 mRNA as a reference control.

### Subcellular fractionation of mouse brain tissues

Subcellular fractionation was performed essentially as described for the rat liver [59] with the following modifications. Freshly harvested brains of 3 mice were homogenised in 3 volumes of 10 mM TrisHCl buffer (pH 7.4) containing 250 mM sucrose, 1 mM EDTA and a full protease inhibitor cocktail using a Potter-Elvehjem homogenizer (30 strokes, on ice). Nuclei and cell debris were removed by a 10-min centrifugation at 800 *g*. Post-nuclear supernatant was further centrifuged for 30 mins using a SW41-Ti Beckman rotor at 10,000 *g* to collect the organellar pellet (“light mitochondrial fraction”) containing lysosomes. The supernatant was further centrifuged using the same rotor for 2 h at 50000 *g* to separate microsomal fraction (pellet) and cytosol. Aliquots of the post-nuclear supernatant, organellar pellet and cytosol were analysed by immunoblot as described above to determine the CTSB content.

### Behavioral analysis

The open-field test was performed as previously described [60]. Briefly, mice were habituated in the experimental room for 30 mins before the commencement of the test.

Each mouse was, then, gently placed in the center of the open-field arena and allowed to explore for 20 min. The mouse was removed and transferred to its home cage after the test, and the arena was cleaned with 70% ethanol before the commencement of the next test. Analysis of the behavioral activity was done using the Smart video tracking software (v3.0, Panlab Harvard Apparatus); the total distance traveled, and percent of time spent in the center zone were measured for hyperactivity and anxiety assessment, respectively.

### E64 treatment

Male and female 8-10-week-old C57BL/6 WT, *Hgsnat^P304L^* and *Neu1^ΔEx3^* mice were randomly assigned to either treatment (E64) or control (vehicle) groups each containing 4-5 male and female mice. Mice in the treatment group received daily intranasal administration of E64 (Sigma-Aldrich, St. Louis, MO, USA, Cat.# E3132) at a dose of 1 mg/kg BW dissolved in saline (0.9% sodium chloride). *Hgsnat^P304L^* mice and their corresponding WT controls were treated for a duration of one month between 5 and 6 months of age. *Neu1^ΔEx3^* mice received the treatment for a duration of 2.5 months between the age of 6 weeks and 4 months. The mice in the vehicle control groups received daily intranasal administration of saline. At treatment days, mice were gently restrained, and E64 solution or vehicle was administered intranasally using a micro-pipette. The administration volume was 5 µl per nostril, administered alternately. Care was taken to ensure uniformity in the administration technique across all animals. At the end of the one-month treatment period, mice were euthanized, and brain tissues harvested and processed for further analyses.

### Statistical analysis

Statistical analyses were performed using Prism GraphPad 9.3.0. software (GraphPad Software San Diego, CA). The normality for all data was analysed using the D’Agostino & Pearson omnibus test. Significance of the difference was determined using t-test (normal distribution) or Mann-Whitney test, when comparing two groups. One-way ANOVA or Nested ANOVA tests, followed by Tukey’s multiple comparisons test (normal distribution), or Kruskal-Wallis test, followed by Dunn’s multiple comparisons test, were used when comparing more than two groups. Two-way ANOVA followed by Tukey post hoc test was used for two-factor analysis. A P-value of 0.05 or less was considered significant.

### Availability of data and materials

The datasets used and/or analyzed during the current study are available from the corresponding author on reasonable request.

## Acknowledgements

The authors thank Dr. Christopher W. Cairo for a generous gift of fluorescently labeled GALC3 protein. We also are grateful to Elke Küster-Schöck and the Plateforme d’Imagerie Microscopique (PIM – CHU Sainte Justine) for the help with life imaging microscopy, and Dr. Mila Ashmarina for critically reading the manuscript and helpful advice.

This work has been partially supported by operating grants PJT-156345 and PJT-180546 from the Canadian Institutes of Health Research, a research grant ND-1 from the Canadian Glycomics Network, Elisa Linton Research Chair in Lysosomal Diseases and research grant co-funded by Sanfilippo Children’s Foundation and Cure Sanfilippo Foundation to A.V.P. G.M.V. was partially supported by Foreign Internship Scholarship (BEPE) from the Fundação de Amparo à Pesquisa do Estado de São Paulo (FAPESP). Approval for the animal experimentation was granted by the Animal Care and Use Committee of the CHU Sainte-Justine.

## Authors’ contributions

Conducted experiments and acquired data: G.M.V., X.P., S.F., MM.X., A.W.; analyzed data: G.M.V., X.P., S.F., MM.X. and A.V.P.; provided essential resources: A.V.P.; wrote the manuscript (first draft): A.V.P., X.P., G.M.V.; wrote the manuscript (editing): X.P., G.M.V. and A.V.P. All authors read and approved the final manuscript.

---

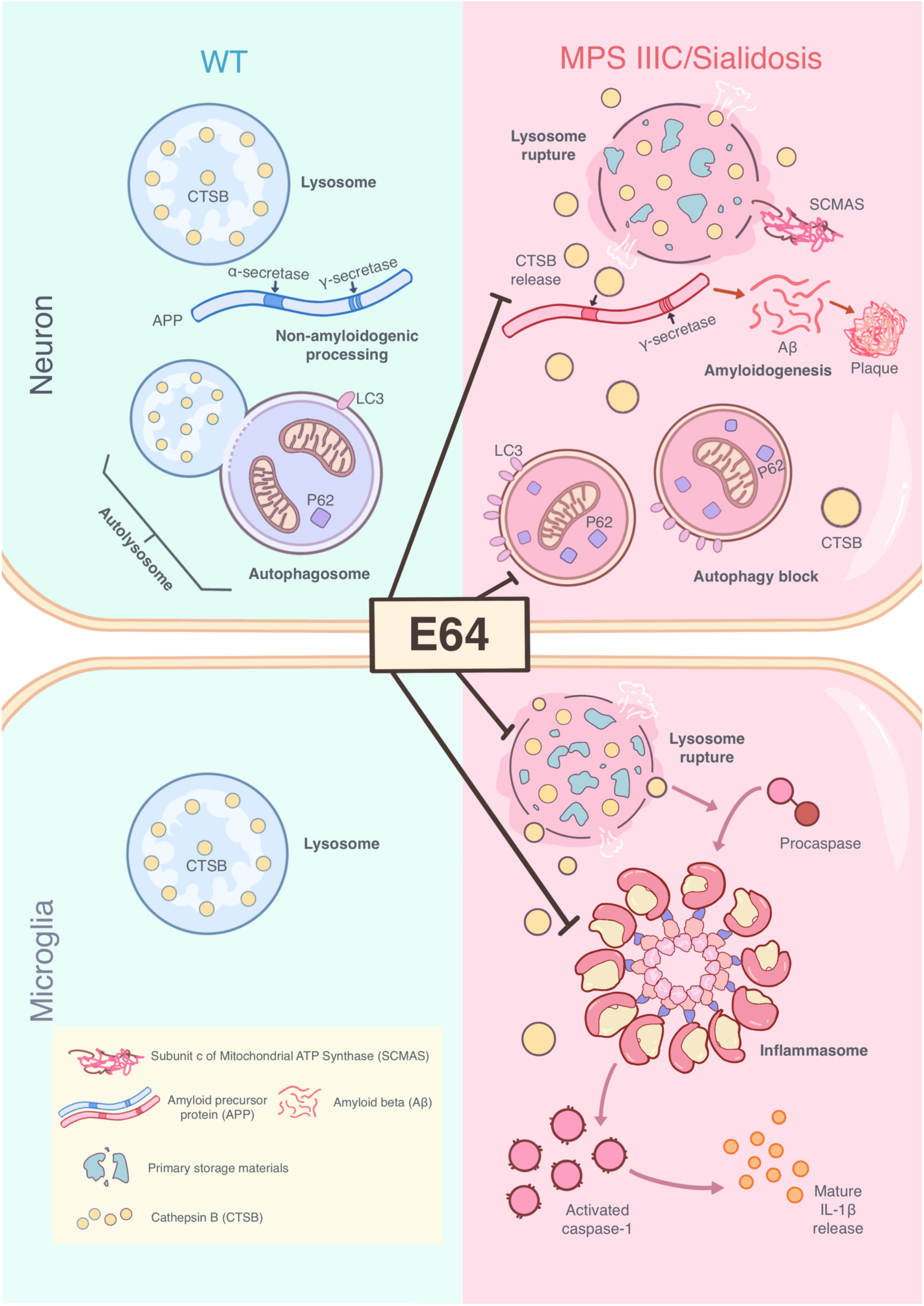

In the healthy neurons of WT mice (left) APP undergoes non-amyloidogenic processing by α– and γ-secretases. In the neurons of MPS IIIC and sialidosis mice (right), CTSB levels are drastically increased and the enzyme partially leaks from lysosomes to the cytoplasm where it participates in the amyloidogenic cleavage of APP. In the microglia, CTSB is an essential component of the inflammasome activation required for secretion of the proinflammatory cytokine, IL-1β. By inhibiting CTSB activity, E64 blocks amyloidogenic cleavage of APP and induces autophagy in MPS IIIC and sialidosis neurons. By inhibiting CTSB in the microglia, the drug reduces release of mature IL-1β.

## References

1. Giugliani, R. Newborn screening for lysosomal diseases: current status and potential interface with population medical genetics in Latin America. J Inherit Metab Dis 2012, 35, 871–877, doi:10.1007/s10545-011-9436-z.

2. Para, C.; Bose, P.; Pshezhetsky, A.V. Neuropathophysiology of Lysosomal Storage Diseases: Synaptic Dysfunction as a Starting Point for Disease Progression. J Clin Med 2020, 9, doi:10.3390/jcm9030616.

3. Ebrahimi-Fakhari, D.; Wahlster, L.; Hoffmann, G.F.; Kolker, S. Emerging role of autophagy in pediatric neurodegenerative and neurometabolic diseases. Pediatric research 2014, 75, 217–226, doi:10.1038/pr.2013.185.

4. Ghavami, S.; Shojaei, S.; Yeganeh, B.; Ande, S.R.; Jangamreddy, J.R.; Mehrpour, M.; Christoffersson, J.; Chaabane, W.; Moghadam, A.R.; Kashani, H.H., et al. Autophagy and apoptosis dysfunction in neurodegenerative disorders. Progress in neurobiology 2014, 112, 24–49, doi:10.1016/j.pneurobio.2013.10.004.

5. Yoon, S.Y.; Kim, D.H. Alzheimer’s disease genes and autophagy. Brain research 2016, 1649, 201–209, doi:10.1016/j.brainres.2016.03.018.

6. Zhang, Y.D.; Zhao, J.J. TFEB Participates in the Abeta-Induced Pathogenesis of Alzheimer’s Disease by Regulating the Autophagy-Lysosome Pathway. DNA and cell biology 2015, 34, 661–668, doi:10.1089/dna.2014.2738.

7. Cho, S.J.; Yun, S.M.; Jo, C.; Lee, D.H.; Choi, K.J.; Song, J.C.; Park, S.I.; Kim, Y.J.; Koh, Y.H. SUMO1 promotes Abeta production via the modulation of autophagy. Autophagy 2015, 11, 100–112, doi:10.4161/15548627.2014.984283.

8. 8. Neufeld, E.F.; Muenzer, J. The mucopolysaccharidoses. In The metabolic basis of inherited disease, 8th ed.; McGraw-Hill, Ed. New York, 2001; pp. 3421-3452.

9. Ohmi, K.; Zhao, H.Z.; Neufeld, E.F. Defects in the medial entorhinal cortex and dentate gyrus in the mouse model of Sanfilippo syndrome type B. PloS one 2011, 6, e27461, doi:10.1371/journal.pone.0027461.

10. Pshezhetsky, A.V. Lysosomal storage of heparan sulfate causes mitochondrial defects, altered autophagy, and neuronal death in the mouse model of mucopolysaccharidosis III type C. Autophagy 2016, 12, 1059–1060, doi:10.1080/15548627.2015.1046671.

11. Beard, H.; Hassiotis, S.; Gai, W.P.; Parkinson-Lawrence, E.; Hopwood, J.J.; Hemsley, K.M. Axonal dystrophy in the brain of mice with Sanfilippo syndrome. Exp Neurol 2017, 295, 243–255, doi:10.1016/j.expneurol.2017.06.010.

12. Wilkinson, F.L.; Holley, R.J.; Langford-Smith, K.J.; Badrinath, S.; Liao, A.; Langford-Smith, A.; Cooper, J.D.; Jones, S.A.; Wraith, J.E.; Wynn, R.F., et al. Neuropathology in mouse models of mucopolysaccharidosis type I, IIIA and IIIB. PloS one 2012, 7, e35787, doi:10.1371/journal.pone.0035787.

13. Martins, C.; Hulkova, H.; Dridi, L.; Dormoy-Raclet, V.; Grigoryeva, L.; Choi, Y.; Langford-Smith, A.; Wilkinson, F.L.; Ohmi, K.; DiCristo, G., et al. Neuroinflammation, mitochondrial defects and neurodegeneration in mucopolysaccharidosis III type C mouse model. Brain 2015, 138, 336–355, doi:10.1093/brain/awu355.

14. Ryazantsev, S.; Yu, W.H.; Zhao, H.Z.; Neufeld, E.F.; Ohmi, K. Lysosomal accumulation of SCMAS (subunit c of mitochondrial ATP synthase) in neurons of the mouse model of mucopolysaccharidosis III B. Molecular genetics and metabolism 2007, 90, 393–401, doi:10.1016/j.ymgme.2006.11.006.

15. Ohmi, K.; Greenberg, D.S.; Rajavel, K.S.; Ryazantsev, S.; Li, H.H.; Neufeld, E.F. Activated microglia in cortex of mouse models of mucopolysaccharidoses I and IIIB. Proc Natl Acad Sci U S A 2003, 100, 1902–1907, doi:10.1073/pnas.252784899.

16. Dawson, G.; Fuller, M.; Helmsley, K.M.; Hopwood, J.J. Abnormal gangliosides are localized in lipid rafts in Sanfilippo (MPS3a) mouse brain. Neurochemical research 2012, 37, 1372–1380, doi:10.1007/s11064-012-0761-x.

17. Pierzynowska, K.; Gaffke, L.; Podlacha, M.; Brokowska, J.; Wegrzyn, G. Mucopolysaccharidosis and Autophagy: Controversies on the Contribution of the Process to the Pathogenesis and Possible Therapeutic Applications. Neuromolecular Med 2020, 22, 25–30, doi:10.1007/s12017-019-08559-1.

18. Scarcella, M.; Scerra, G.; Ciampa, M.; Caterino, M.; Costanzo, M.; Rinaldi, L.; Feliciello, A.; Anzilotti, S.; Fiorentino, C.; Renna, M., et al. Metabolic rewiring and autophagy inhibition correct lysosomal storage disease in mucopolysaccharidosis IIIB. iScience 2024, 27, 108959, doi:10.1016/j.isci.2024.108959.

19. Maeda, M.; Seto, T.; Kadono, C.; Morimoto, H.; Kida, S.; Suga, M.; Nakamura, M.; Kataoka, Y.; Hamazaki, T.; Shintaku, H. Autophagy in the Central Nervous System and Effects of Chloroquine in Mucopolysaccharidosis Type II Mice. Int J Mol Sci 2019, 20, doi:10.3390/ijms20235829.

20. Capuozzo, A.; Montefusco, S.; Cacace, V.; Sofia, M.; Esposito, A.; Napolitano, G.; Nusco, E.; Polishchuk, E.; Pizzo, M.T.; De Risi, M., et al. Fluoxetine ameliorates mucopolysaccharidosis type IIIA. Mol Ther 2022, 30, 1432–1450, doi:10.1016/j.ymthe.2022.01.037.

21. Annunziata, I.; Patterson, A.; Helton, D.; Hu, H.; Moshiach, S.; Gomero, E.; Nixon, R.; d’Azzo, A. Lysosomal NEU1 deficiency affects amyloid precursor protein levels and amyloid-beta secretion via deregulated lysosomal exocytosis. Nat Commun 2013, 4, 2734, doi:10.1038/ncomms3734.

22. Haass, C.; Kaether, C.; Thinakaran, G.; Sisodia, S. Trafficking and proteolytic processing of APP. Cold Spring Harbor perspectives in medicine 2012, 2, a006270, doi:10.1101/cshperspect.a006270.

23. Li, R.; Lindholm, K.; Yang, L.B.; Yue, X.; Citron, M.; Yan, R.; Beach, T.; Sue, L.; Sabbagh, M.; Cai, H., et al. Amyloid beta peptide load is correlated with increased beta-secretase activity in sporadic Alzheimer’s disease patients. Proc Natl Acad Sci U S A 2004, 101, 3632–3637, doi:10.1073/pnas.0205689101.

24. Fukumoto, H.; Cheung, B.S.; Hyman, B.T.; Irizarry, M.C. Beta-secretase protein and activity are increased in the neocortex in Alzheimer disease. Archives of neurology 2002, 59, 1381–1389.

25. Hook, V.; Schechter, I.; Demuth, H.U.; Hook, G. Alternative pathways for production of beta-amyloid peptides of Alzheimer’s disease. Biological chemistry 2008, 389, 993–1006, doi:10.1515/BC.2008.124.

26. Hook, V.; Toneff, T.; Bogyo, M.; Greenbaum, D.; Medzihradszky, K.F.; Neveu, J.; Lane, W.; Hook, G.; Reisine, T. Inhibition of cathepsin B reduces beta-amyloid production in regulated secretory vesicles of neuronal chromaffin cells: evidence for cathepsin B as a candidate beta-secretase of Alzheimer’s disease. Biological chemistry 2005, 386, 931–940, doi:10.1515/BC.2005.108.

27. Chevallier, N.; Vizzavona, J.; Marambaud, P.; Baur, C.P.; Spillantini, M.; Fulcrand, P.; Martinez, J.; Goedert, M.; Vincent, J.P.; Checler, F. Cathepsin D displays in vitro beta-secretase-like specificity. Brain research 1997, 750, 11–19.

28. Di Domenico, F.; Coccia, R.; Cocciolo, A.; Murphy, M.P.; Cenini, G.; Head, E.; Butterfield, D.A.; Giorgi, A.; Schinina, M.E.; Mancuso, C., et al. Impairment of proteostasis network in Down syndrome prior to the development of Alzheimer’s disease neuropathology: redox proteomics analysis of human brain. Biochimica et biophysica acta 2013, 1832, 1249–1259, doi:10.1016/j.bbadis.2013.04.013.

29. Sun, B.; Zhou, Y.; Halabisky, B.; Lo, I.; Cho, S.H.; Mueller-Steiner, S.; Devidze, N.; Wang, X.; Grubb, A.; Gan, L. Cystatin C-cathepsin B axis regulates amyloid beta levels and associated neuronal deficits in an animal model of Alzheimer’s disease. Neuron 2008, 60, 247–257, doi:10.1016/j.neuron.2008.10.001.

30. Wu, Z.; Sun, L.; Hashioka, S.; Yu, S.; Schwab, C.; Okada, R.; Hayashi, Y.; McGeer, P.L.; Nakanishi, H. Differential pathways for interleukin-1beta production activated by chromogranin A and amyloid beta in microglia. Neurobiol Aging 2013, 34, 2715–2725, doi:10.1016/j.neurobiolaging.2013.05.018.

31. Qiao, C.; Yin, N.; Gu, H.Y.; Zhu, J.L.; Ding, J.H.; Lu, M.; Hu, G. Atp13a2 Deficiency Aggravates Astrocyte-Mediated Neuroinflammation via NLRP3 Inflammasome Activation. CNS Neurosci Ther 2016, 22, 451–460, doi:10.1111/cns.12514.

32. Ni, J.; Wu, Z.; Stoka, V.; Meng, J.; Hayashi, Y.; Peters, C.; Qing, H.; Turk, V.; Nakanishi, H. Increased expression and altered subcellular distribution of cathepsin B in microglia induce cognitive impairment through oxidative stress and inflammatory response in mice. Aging Cell 2019, 18, e12856, doi:10.1111/acel.12856.

33. Man, S.M.; Kanneganti, T.D. Regulation of lysosomal dynamics and autophagy by CTSB/cathepsin B. Autophagy 2016, 12, 2504–2505, doi:10.1080/15548627.2016.1239679.

34. Viana, G.M.; Gonzalez, E.A.; Alvarez, M.M.P.; Cavalheiro, R.P.; do Nascimento, C.C.; Baldo, G.; D’Almeida, V.; de Lima, M.A.; Pshezhetsky, A.V.; Nader, H.B. Cathepsin B-associated Activation of Amyloidogenic Pathway in Murine Mucopolysaccharidosis Type I Brain Cortex. Int J Mol Sci 2020, 21, doi:10.3390/ijms21041459.

35. Pan, X.; Taherzadeh, M.; Bose, P.; Heon-Roberts, R.; Nguyen, A.L.A.; Xu, T.; Para, C.; Yamanaka, Y.; Priestman, D.A.; Platt, F.M., et al. Glucosamine amends CNS pathology in mucopolysaccharidosis IIIC mouse expressing misfolded HGSNAT. J Exp Med 2022, 219, doi:10.1084/jem.20211860.

36. Kho, I.; Demina, E.P.; Pan, X.; Londono, I.; Cairo, C.W.; Sturiale, L.; Palmigiano, A.; Messina, A.; Garozzo, D.; Ung, R.V., et al. Severe kidney dysfunction in sialidosis mice reveals an essential role for neuraminidase 1 in reabsorption. JCI Insight 2023, 8, doi:10.1172/jci.insight.166470.

37. Lowden, J.A.; O’Brien, J.S. Sialidosis: a review of human neuraminidase deficiency. Am J Hum Genet 1979, 31, 1–18.

38. Pshezhetsky, A.V.; Richard, C.; Michaud, L.; Igdoura, S.; Wang, S.; Elsliger, M.A.; Qu, J.; Leclerc, D.; Gravel, R.; Dallaire, L., et al. Cloning, expression and chromosomal mapping of human lysosomal sialidase and characterization of mutations in sialidosis. Nat Genet 1997, 15, 316–320, doi:10.1038/ng0397-316.

39. Yang, E.H.; Rode, J.; Howlader, M.A.; Eckermann, M.; Santos, J.T.; Hernandez Armada, D.; Zheng, R.; Zou, C.; Cairo, C.W. Galectin-3 alters the lateral mobility and clustering of beta1-integrin receptors. PLoS One 2017, 12, e0184378, doi:10.1371/journal.pone.0184378.

40. Hook, V.Y.; Kindy, M.; Hook, G. Inhibitors of cathepsin B improve memory and reduce beta-amyloid in transgenic Alzheimer disease mice expressing the wild-type, but not the Swedish mutant, beta-secretase site of the amyloid precursor protein. J Biol Chem 2008, 283, 7745–7753, doi:10.1074/jbc.M708362200.

41. Hook, V.; Hook, G.; Kindy, M. Pharmacogenetic features of cathepsin B inhibitors that improve memory deficit and reduce beta-amyloid related to Alzheimer’s disease. Biol Chem 2010, 391, 861–872, doi:10.1515/BC.2010.110.

42. Hook, G.; Jacobsen, J.S.; Grabstein, K.; Kindy, M.; Hook, V. Cathepsin B is a New Drug Target for Traumatic Brain Injury Therapeutics: Evidence for E64d as a Promising Lead Drug Candidate. Front Neurol 2015, 6, 178, doi:10.3389/fneur.2015.00178.

43. Hook, G.; Hook, V.; Kindy, M. The cysteine protease inhibitor, E64d, reduces brain amyloid-beta and improves memory deficits in Alzheimer’s disease animal models by inhibiting cathepsin B, but not BACE1, beta-secretase activity. J Alzheimers Dis 2011, *26*, 387-408, doi:10.3233/JAD-2011-110101.

44. Dhuyvetter, D.; Tekle, F.; Nazarov, M.; Vreeken, R.J.; Borghys, H.; Rombouts, F.; Lenaerts, I.; Bottelbergs, A. Direct nose to brain delivery of small molecules: critical analysis of data from a standardized in vivo screening model in rats. Drug Deliv 2020, 27, 1597–1607, doi:10.1080/10717544.2020.1837291.

45. Biffi, A.; Montini, E.; Lorioli, L.; Cesani, M.; Fumagalli, F.; Plati, T.; Baldoli, C.; Martino, S.; Calabria, A.; Canale, S., et al. Lentiviral hematopoietic stem cell gene therapy benefits metachromatic leukodystrophy. Science 2013, 341, 1233158, doi:10.1126/science.1233158.

46. Sessa, M.; Lorioli, L.; Fumagalli, F.; Acquati, S.; Redaelli, D.; Baldoli, C.; Canale, S.; Lopez, I.D.; Morena, F.; Calabria, A., et al. Lentiviral haemopoietic stem-cell gene therapy in early-onset metachromatic leukodystrophy: an ad-hoc analysis of a non-randomised, open-label, phase 1/2 trial. Lancet 2016, 388, 476–487, doi:10.1016/S0140-6736(16)30374-9.

47. Hook, G.; Reinheckel, T.; Ni, J.; Wu, Z.; Kindy, M.; Peters, C.; Hook, V. Cathepsin B Gene Knockout Improves Behavioral Deficits and Reduces Pathology in Models of Neurologic Disorders. Pharmacol Rev 2022, 74, 600–629, doi:10.1124/pharmrev.121.000527.

48. Moon, H.Y.; Becke, A.; Berron, D.; Becker, B.; Sah, N.; Benoni, G.; Janke, E.; Lubejko, S.T.; Greig, N.H.; Mattison, J.A., et al. Running-Induced Systemic Cathepsin B Secretion Is Associated with Memory Function. Cell Metab 2016, 24, 332–340, doi:10.1016/j.cmet.2016.05.025.

49. Turk, V.; Stoka, V.; Vasiljeva, O.; Renko, M.; Sun, T.; Turk, B.; Turk, D. Cysteine cathepsins: from structure, function and regulation to new frontiers. Biochim Biophys Acta 2012, 1824, 68–88, doi:10.1016/j.bbapap.2011.10.002.

50. Yoon, M.C.; Hook, V.; O’Donoghue, A.J. Cathepsin B Dipeptidyl Carboxypeptidase and Endopeptidase Activities Demonstrated across a Broad pH Range. Biochemistry 2022, 61, 1904–1914, doi:10.1021/acs.biochem.2c00358.

51. Pan, X.; Taherzadeh, M.; Bose, P.; Heon-Roberts, R.; Nguyen, A.L.; Xu, T.; Pará, C.; Yamanaka, Y.; Priestman, D.A.; Platt, F.M. Glucosamine amends CNS pathology in mucopolysaccharidosis IIIC mouse expressing misfolded HGSNAT. Journal of Experimental Medicine 2022, 219, e20211860.

52. Jones-Tabah, J.; He, K.; Senkevich, K.; Karpilovsky, N.; Deyab, G.; Cousineau, Y.; Nikanorova, D.; Goldsmith, T.; Del Cid Pellitero, E.; Chen, C.X., et al. The Parkinson’s disease risk gene cathepsin B promotes fibrillar alpha-synuclein clearance, lysosomal function and glucocerebrosidase activity in dopaminergic neurons. *bioRxiv* 2023, 10.1101/2023.11.11.566693, doi:10.1101/2023.11.11.566693.

53. Ni, J.; Lan, F.; Xu, Y.; Nakanishi, H.; Li, X. Extralysosomal cathepsin B in central nervous system: Mechanisms and therapeutic implications. Brain Pathol 2022, 32, e13071, doi:10.1111/bpa.13071.

54. Polgreen, L.E.; Chen, A.H.; Pak, Y.; Luzzi, A.; Morales Garval, A.; Acevedo, J.; Bitan, G.; Iacovino, M.; O’Neill, C.; Eisengart, J.B. Author Correction: Anakinra in Sanfilippo syndrome: a phase 1/2 trial. Nat Med 2024, 30, 2693, doi:10.1038/s41591-024-03207-z.

55. Luo, C.L.; Chen, X.P.; Yang, R.; Sun, Y.X.; Li, Q.Q.; Bao, H.J.; Cao, Q.Q.; Ni, H.; Qin, Z.H.; Tao, L.Y. Cathepsin B contributes to traumatic brain injury-induced cell death through a mitochondria-mediated apoptotic pathway. J Neurosci Res 2010, 88, 2847–2858, doi:10.1002/jnr.22453.

56. Rashid Khan, M.; Fayaz Ahmad, S.; Nadeem, A.; Imam, F.; Al-Harbi, N.O.; Shahnawaz Khan, M.; Alsahli, M.; Alhosaini, K. Cathepsin-B inhibitor CA-074 attenuates retinopathy and optic neuritis in experimental autoimmune encephalomyelitis induced in SJL/J mice. Saudi Pharm J 2023, 31, 147–153, doi:10.1016/j.jsps.2022.11.013.

57. Zenon-Melendez, C.N.; Carrasquillo Carrion, K.; Cantres Rosario, Y.; Roche Lima, A.; Melendez, L.M. Inhibition of Cathepsin B and SAPC Secreted by HIV-Infected Macrophages Reverses Common and Unique Apoptosis Pathways. J Proteome Res 2022, 21, 301–312, doi:10.1021/acs.jproteome.1c00187.

58. Pan, X.; De Aragao, C.B.P.; Velasco-Martin, J.P.; Priestman, D.A.; Wu, H.Y.; Takahashi, K.; Yamaguchi, K.; Sturiale, L.; Garozzo, D.; Platt, F.M., et al. Neuraminidases 3 and 4 regulate neuronal function by catabolizing brain gangliosides. FASEB J 2017, 31, 3467–3483, doi:10.1096/fj.201601299R.

59. 59. Evans, W.H. Isolation and characterization of membranes and cell organelles. In PreparativeCentrifugation - A Practical Approach, Rickwood, D., Ed. Oxford University Press: Oxford, UK, 1992; pp. 233–270.

60. Amegandjin, C.A.; Choudhury, M.; Jadhav, V.; Carrico, J.N.; Quintal, A.; Berryer, M.; Snapyan, M.; Chattopadhyaya, B.; Saghatelyan, A.; Di Cristo, G. Sensitive period for rescuing parvalbumin interneurons connectivity and social behavior deficits caused by TSC1 loss. Nat Commun 2021, 12, 3653, doi:10.1038/s41467-021-23939-7.

