## Supplementary materials for "Inhibition of cathepsin B blocks amyloidogenesis in the mouse models of neurological lysosomal diseases mucopolysaccharidosis type IIIC and sialidosis"

**Table S1. Antibodies and their working concentrations used in the current work.**

| Antigen | Host/Target species | Dilution | Manufacturer |
|---|---|---|---|
| GFAP | Rabbit anti-mouse | 1:300 | DSHB (8-1E7-s) |
| Lysosome-associated membrane protein 2 (LAMP2) | Rat anti-mouse | 1:200 | DSHB (ABL-93-s) |
| NeuN | Rabbit anti-mouse | 1:200 | Millipore Sigma (MABN140) |
| CD68 | Rabbit polyclonal to CD68 | 1:200 | Abcam (ab125212) |
| $G_{M2}$-ganglioside | Mouse humanized | 1:400 | KM966 |
| β-Amyloid (D54D2)<br>CD11b | Rabbit anti-mouse<br>Rat anti-mouse | 1:200<br>1:50 | Cell Signaling (8243S)<br>DHSB (M1/70.15.11.5.2) |
| P62 | Mouse | 1:200 | BD BIOSCIENCES (610832) |

| | | | |
|---|---|---|---|
| LC3b | Rabbit | 1:200 | GeneTex (GTX82986) |
| APP Antibody | Rabbit | 1:200 | Cell Signaling (2452) |
| Cathepsin B (D1C7Y) XP® Rabbit mAb | Rabbit | 1:200 | Cell Signaling (31718S) |
| Recombinant Anti-ATP synthase C antibody (SCMAS) | Rabbit anti-mouse | 1:200 | Abcam (ab181243) |
| IgG | Goat anti-rabbit, anti-mouse, or anti-rat Alexa Fluor 488-, Alexa Fluor 555- or Alexa Fluor 633-conjugated | 1:400 | Thermo Fisher Scientific |

**Table S2. Forward (F) and reverse (R) primers used for real-time qPCR.**

| **Gene** | **Sequence** |
| --- | --- |
| F-IL-1β | TGAAATGCCACCTTTTGACA |
| R-IL-1β | GTAGCTGCCACAGCTTCTCC |
| F-MIP1α | GCCCTTGCTGTTCTTCTCTG |
| R-MIP1α | CAGATCTGCCGGTTTCTCTT |
| F-TNFα | TCTTCTCATTCCTGCTTGTGG |
| R-TNFα | CACTTGGTGGTTTGCTACGA |
| F-RPL32 | TTCTTCCTCGGCGCTGCCTACGA |
| R-RPL32 | AACCTTCTCCGCACCCTGTTGTCA |

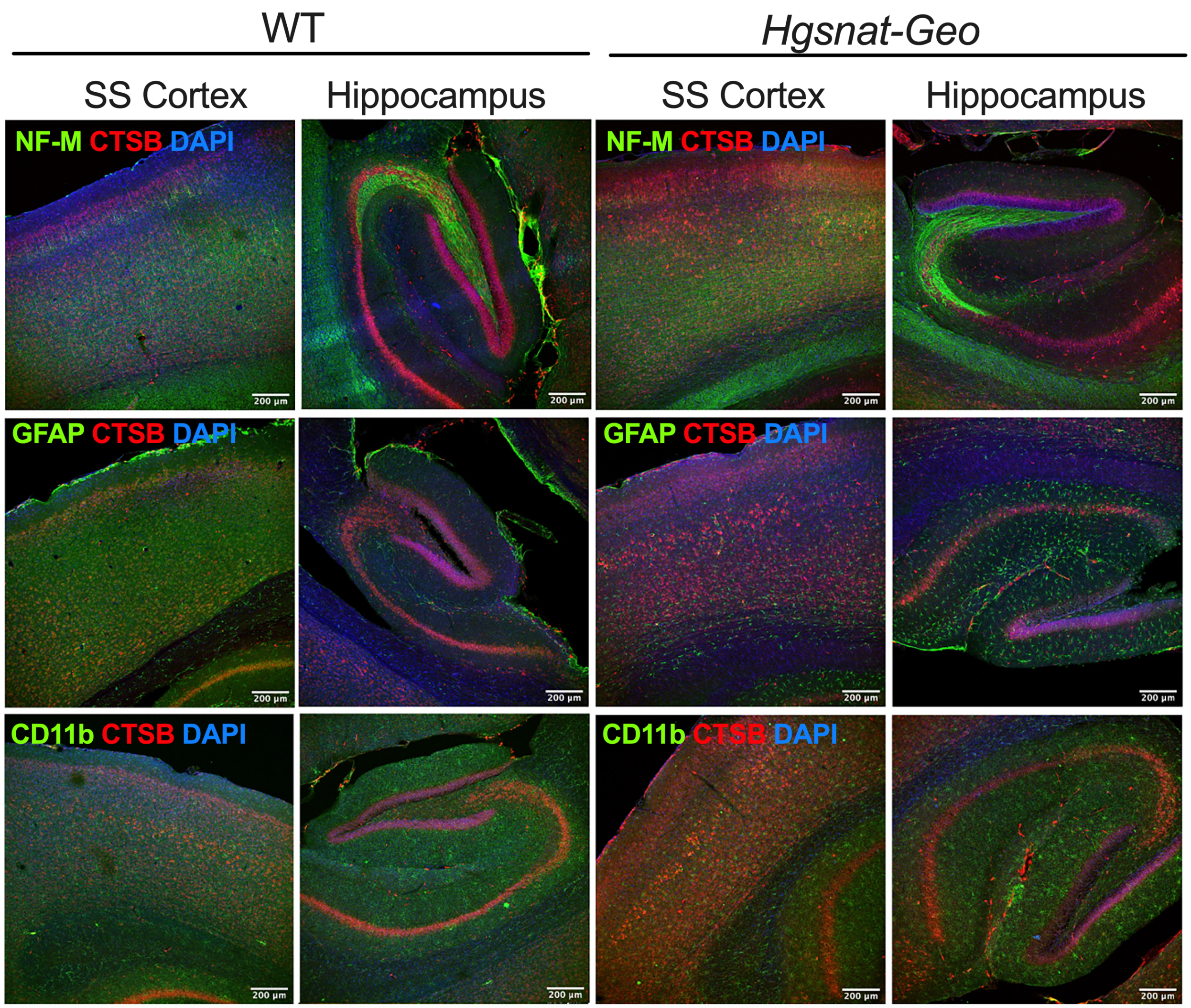


**Figure S1. CTSB levels are increased in cortical but not in hippocampal neurons of *Hgsnat-Geo* compared to WT mice.**

Panels show representative confocal microscopy images of SS cortices and hippocampi of 6-month-old WT and *Hgsnat-Geo* mice labeled for CTSB (red) and Neurofilament medium chain (NF-M, green), GFAP (green), or CD11b (green). DAPI (blue) was used as a nuclear counterstain. Scale bars equal 200 μm.

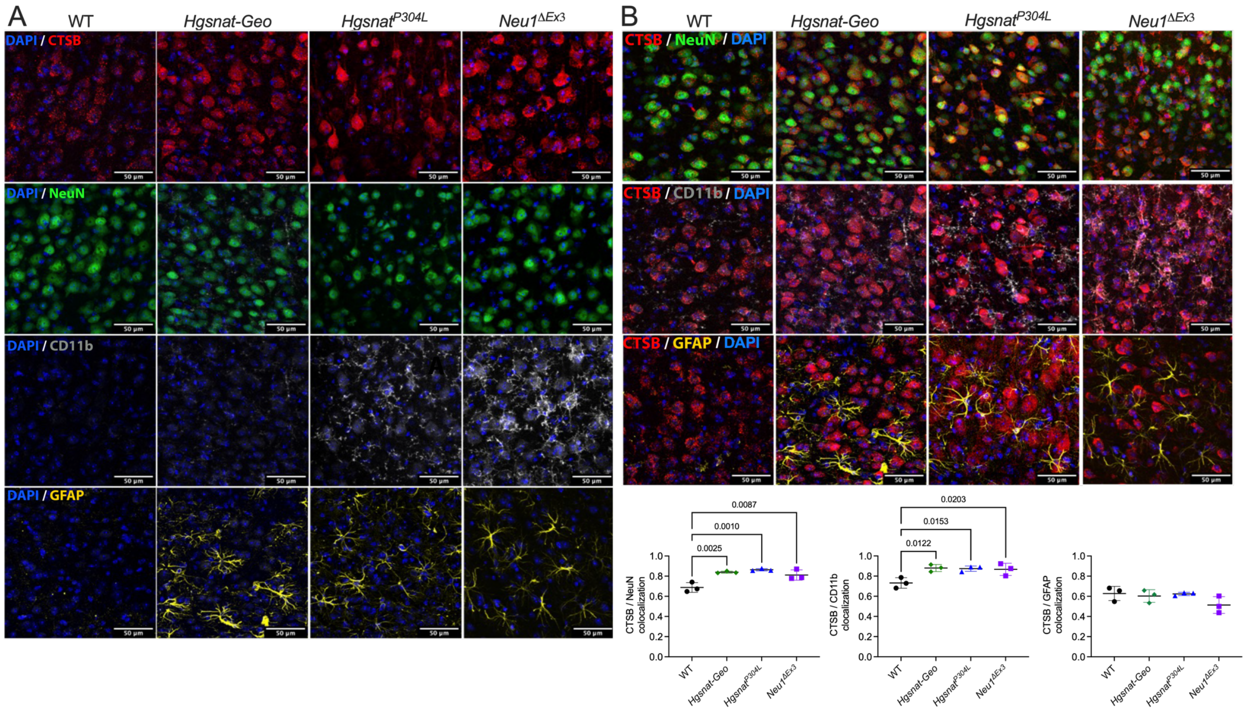


**Figure S2. CTSB colocalizes with NeuN-positive neurons and CD11b-positive microglia but not with GFAP-positive astrocytes in brain cortices of MPS IIIC and sialidosis mice.**

**(A)** Representative confocal microscopy images of brain cortex (layer V) of WT, *Hgsnat-Geo*, *Hgsnat*$^{P304L}$ and *Neu1*$^{\Delta Ex3}$ mice showing immunostaining for CTSB (red), cortical neurons (NeuN, green), activated microglia (CD11b, gray) and astrocytes (GFAP, yellow). **(B)** Representative images and quantitative analysis (Manders' overlap coefficient) of CATB/NeuN, CATB/CD11b and CATB/GFAP colocalization. Graphs show individual data, means and SD. P values were calculated by ANOVA with Tukey post hoc test; n = 3 animals per genotype, thirty images were analyzed for each cortex. Scale bars equal 50 µm.

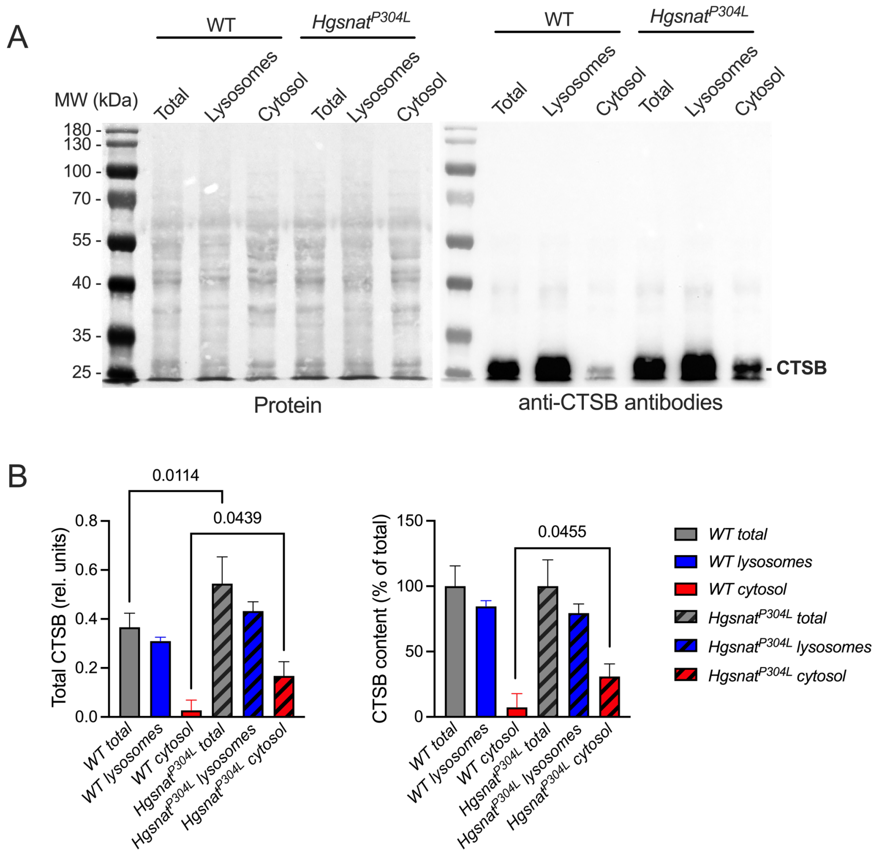


**Figure S3. CTSB levels are increased in the cytosol fraction purified from brain tissues of *Hgsnat*$^{P304L}$ mice compared to that WT mice.**

**(A)** Representative blot images of post-nuclear supernatant (Total), organellar fraction (Lysosome) and cytosol (Cytosol) from pooled tissues of 6-months-old WT or *Hgsnat*$^{P304L}$ mice stained with Ponceau (Protein) or anti-CTSB antibodies. Thirty µg of total protein was loaded on each lane. **(B)** Estimated total (left) and relative (right) amount of CTSB in each fraction. Data show means and SD of 3 independent experiments each time performed with pooled brains of 2 mice per genotype. P values were calculated by one-way ANOVA with Šídák post hoc test.

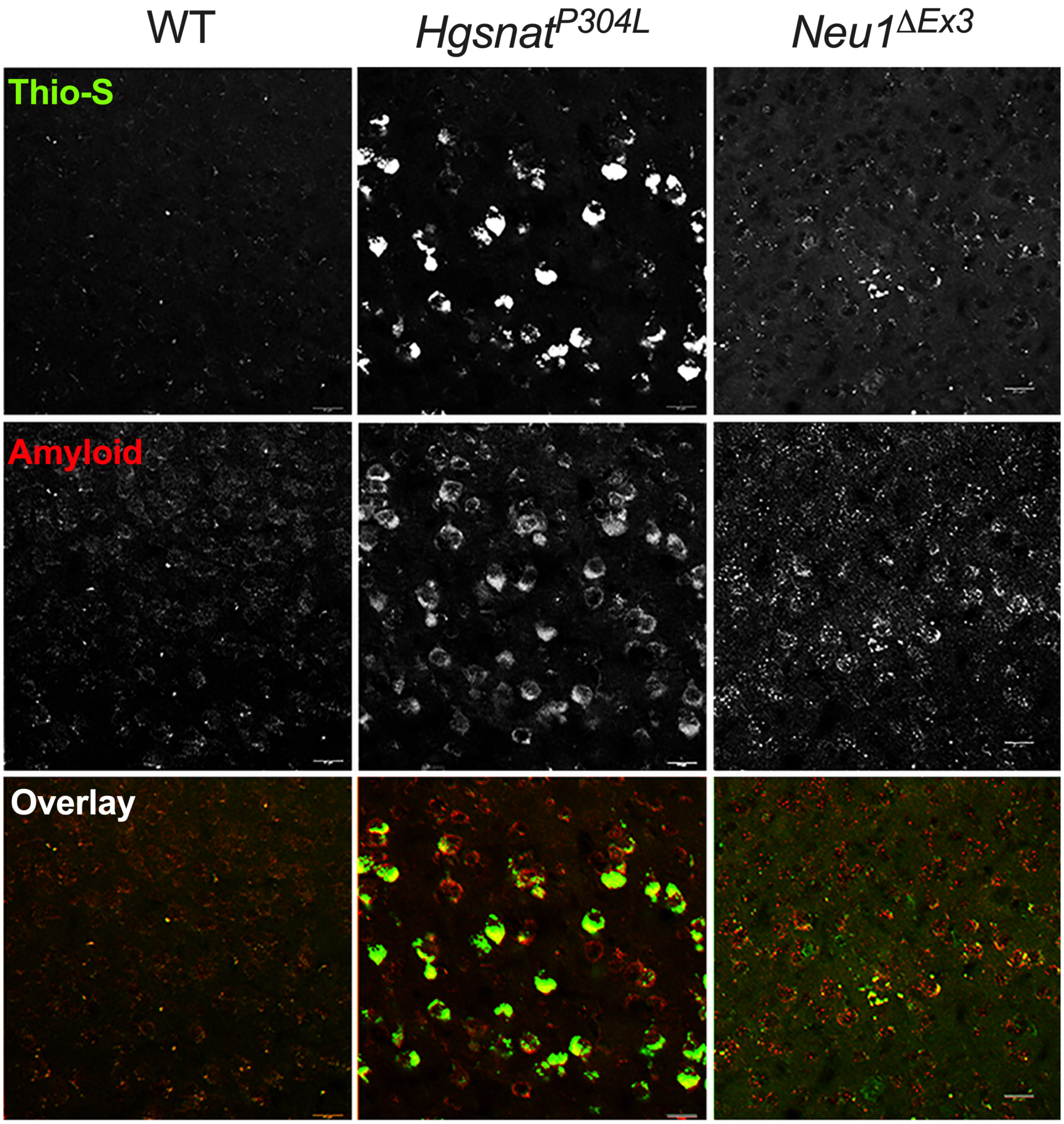


**Figure S4. Pyramidal cortical layers IV-V neurons in the brains of 6-month-old *Hgsnat*$^{P304L}$ and *Neu1*$^{\Delta Ex3}$ but not WT mice show positive labeling for both Thioflavin-S and β-amyloid.**

Panels show representative confocal microscopy images of brain cortices (layers IV-V) of WT, *Hgsnat*$^{P304L}$, and *Neu1*$^{\Delta Ex3}$ mice labeled with Thioflavin-S (Thio-S, green) and antibodies against β-amyloid protein (Amyloid, red). Scale bars equal 25 µm.

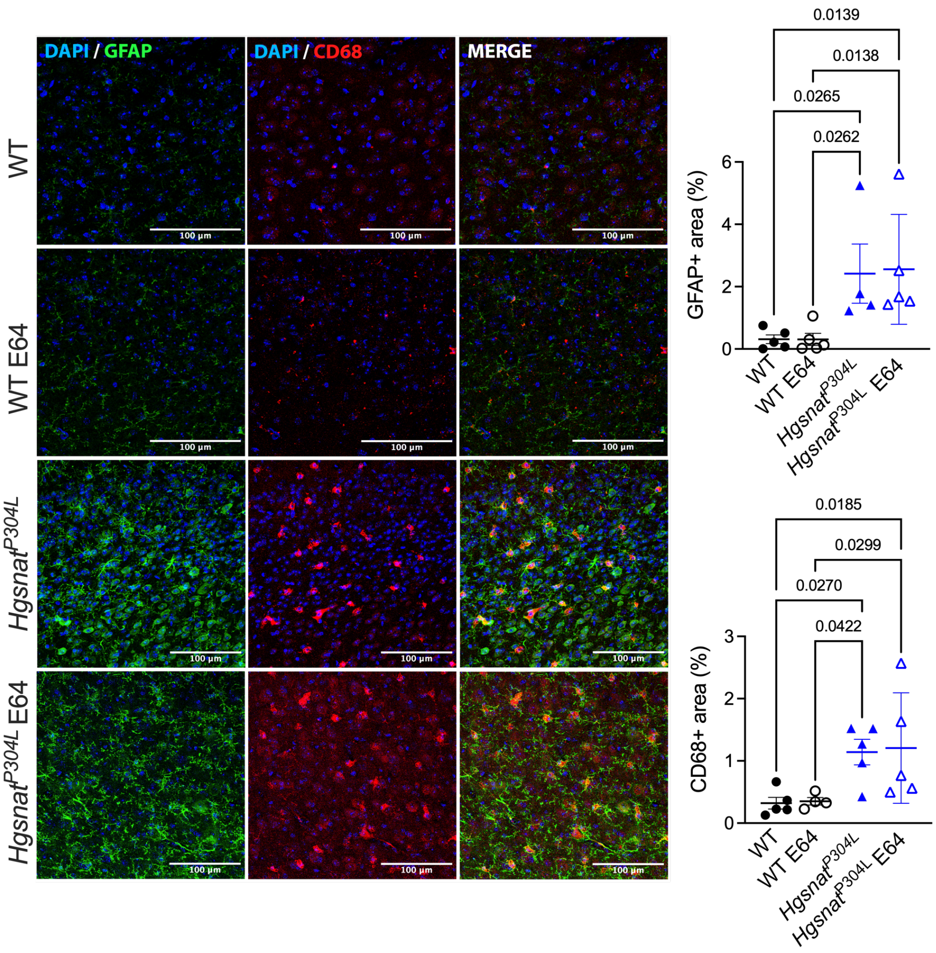


**Figure S5. Six-month-old untreated and E64-treated *Hgsnat*$^{P304L}$ mice show increased microastrogliosis compared to untreated or E64-treated WT mice.**

Panels show representative confocal microscopy images of brain cortices (layers IV-V) of WT, E64-treated WT, *Hgsnat*$^{P304L}$, and E64-treated *Hgsnat*$^{P304L}$ mice labeled for CD68 (red) and GFAP (green). Nuclei were counterstained with DAPI. Scale bars equal 100 µm. Graphs show quantification of immunofluorescence with ImageJ software. Individual data, means and SD are shown. Statistical analysis was performed by ANOVA with Tukey post hoc test; n=4-5 mice per genotype.

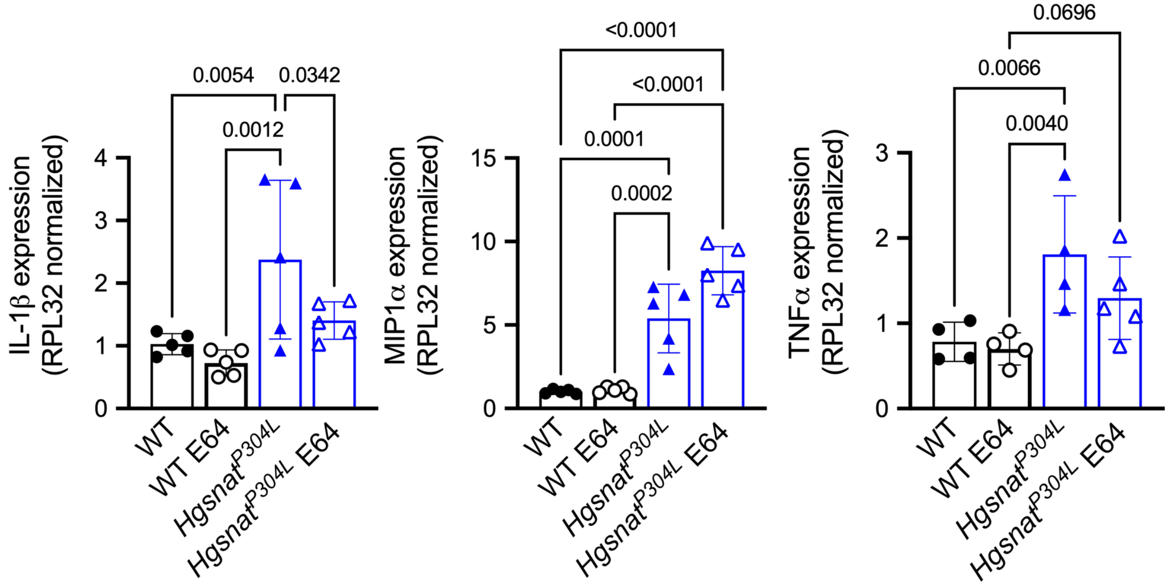


**Figure S6. Six-month-old untreated and E-64 treated *Hgsnat*$^{P304L}$ mice show increased expression of pro-inflammatory cytokines MIP1$\alpha$, and TNF$\alpha$ compared to untreated or E64-treated WT mice.**

Total brain tissues of both untreated and E64-treated *Hgsnat*$^{P304L}$ mice show increased expression of inflammation markers, MIP1$\alpha$, and TNF$\alpha$ compared to untreated or E64-treated WT mice, while expression of IL-1β is ameliorated by the treatment. The cytokine mRNA levels are normalized for the *RLP32* mRNA content. Individual data, means and SD are shown. Statistical analysis was performed by nested ANOVA with Tukey post hoc test; n=5 mice per genotype, two independent experiments were performed with tissues from each mouse. Only p values < 0.07 are shown.
